# Broadband synergy versus oscillatory redundancy in the visual cortex

**DOI:** 10.1101/2025.02.27.640666

**Authors:** Louis Roberts, Juho Äijälä, Florian Burger, Cem Uran, Michael A. Jensen, Kai J. Miller, Robin A.A. Ince, Martin Vinck, Dora Hermes, Andres Canales-Johnson

## Abstract

The cortex generates diverse neural dynamics, ranging from broadband fluctuations to narrowband oscillations at specific frequencies. Here, we investigated whether broadband and oscillatory dynamics play different roles in the encoding and transmission of visual information. We used information-theoretical measures to dissociate neural signals sharing common information (i.e., redundancy) from signals encoding complementary information (i.e., synergy). We analyzed electrocorticography (ECoG) and local field potentials (LFP) in the visual cortex of human and non-human primates (macaque) to investigate the extent to which broadband signals (BB) and narrowband gamma (NBG) oscillations conveyed synergistic or redundant information about images. In both species, the information conveyed by BB signals was highly synergistic within and between visual areas. By contrast, the information carried by NBG was primarily redundant within and between the same visual areas. Finally, the information conveyed by BB signals emerged early after stimulus onset, while NBG sustained information at later time points. These results suggest a potential dual role of BB and NBG cortical dynamics in visual processing, with broadband dynamics supporting nonlinear pattern recognition and oscillations facilitating information maintenance across the cortex.

## INTRODUCTION

The cortex generates a diverse spectrum of dynamics, ranging from broadband activity to oscillatory activity in specific frequency bands (Bressler and Kelso, 2001; Buzsáki, 2006; Freeman and Skarda, 1985; Hermes et al., 2015a; Miller et al., 2009b; Ray and Maunsell, 2011; Varela et al., 2001; Vinck et al., 2023). A major open question is whether interactions between brain areas are mainly based on broadband dynamics or oscillatory interactions. Several theories have emphasized that coordinated interactions are based on narrowband synchronization or resonance phenomena (Bressler and Kelso, 2001; Fries, 2005; Varela et al., 2001; Vinck et al., 2013). In contrast, other theories emphasize the importance of broadband transients and nonlinear relationships across a broad range of frequencies (Sani et al., 2024; Terada and Toyoizumi, 2024; Vinck et al., 2025, 2023; Yang et al., 2021). Thus, it remains an open question what role these dynamics play in communication and whether they play distinct and complementary roles in encoding and transmitting information (Vinck et al., 2023).

An increasing body of work suggests that information about behavioral and sensory variables can be decoded from many locations, highlighting the distributed nature of neural representations (Allen et al., 2019; Chen et al., 2024; Khilkevich et al., 2024; de Schotten and Forkel, 2022; Shenoy and Kao, 2021; Steinmetz et al., 2019; Stringer et al., 2019; Urai et al., 2022). However, distributed representations do not necessarily encode redundant information; they might also encode synergistic information, concepts derived from information theory. In information theory applied to neural signals (Blume et al., 2026; Chidichimo et al., 2025; Combrisson et al., 2024; Daube et al., 2019; Delis et al., 2022; Francis et al., 2022; Gelens et al., 2024; Greco et al., 2024; Ince et al., 2017, 2016; Koçillari et al., 2023; Luppi et al., 2022; Nigam et al., 2019; Olivares et al., 2025; Park et al., 2018; Varley et al., 2023), redundancy measures the shared (or common) information that a set of signals encodes about a stimulus variable. On the other hand, the measure of synergy quantifies the complementary (or extra) information that a set of signals encodes as a whole, above and beyond the information encoded by the sum of the isolated signals. Therefore, redundant and synergistic representations are likely functionally relevant, with redundant representations ensuring robustness but synergistic representations reflecting emergent dynamics in a nonlinear, recurrent coupled system (Gelens et al., 2024). The readout of redundant and synergistic information likely requires different readout mechanisms and network architectures. If their oscillatory or broadband nature governs the integration of distributed signals, then it is plausible that broadband and oscillatory dynamics play different roles in the encoding and transmitting synergistic and redundant information. Previous work has suggested that broadband activity represents auditory prediction errors synergistically in the auditory cortex (Gelens et al., 2024). However, since the activity in the auditory system is dominated by broadband fluctuations rather than oscillatory dynamics (Canales-Johnson et al., 2021), a direct contrast between broadband vs. oscillatory dynamics is still lacking.

Although narrowband oscillatory activity is not prominent in the auditory cortex, broadband fluctuations and narrowband oscillations are commonly superimposed in other model systems such as the visual system (Hermes et al., 2015a,b, 2017, 2019; Ray and Maunsell, 2011). In fact, several theories have proposed a critical functional role for narrowband gamma oscillations in inter-areal communication and visual coding (Aizenbud et al., 2025; Fries, 2005; Fries et al., 2007; Ray and Maunsell, 2015; Salinas and Sejnowski, 2001; Singer, 1999; Vinck and Bosman, 2016; Vinck et al., 2010, 2025, 2013). Therefore, the visual model system is well-suited to test whether broadband and narrow-band activity subserve the encoding of synergistic or redundant information across nodes. Here, we analyzed electrocorticography (ECoG) and local field potentials (LFP) signals in the visual cortex of humans (subjects S1 and S2) and non-human primates (macaque M1) and decomposed signals into broadband and narrowband activity. We then investigated to what extent different electrodes (within and between areas) conveyed redundant or synergistic information about visual images via broadband or narrowband gamma oscillations.

## RESULTS

To understand how mesoscale field potentials in the visual cortex encode and communicate information about image statistics, we analyzed ECoG data from two human subjects (areas V1, V2, and V3) and LFP data from one macaque (areas V1 and V4) (Figure 1A).

**Figure 1:**
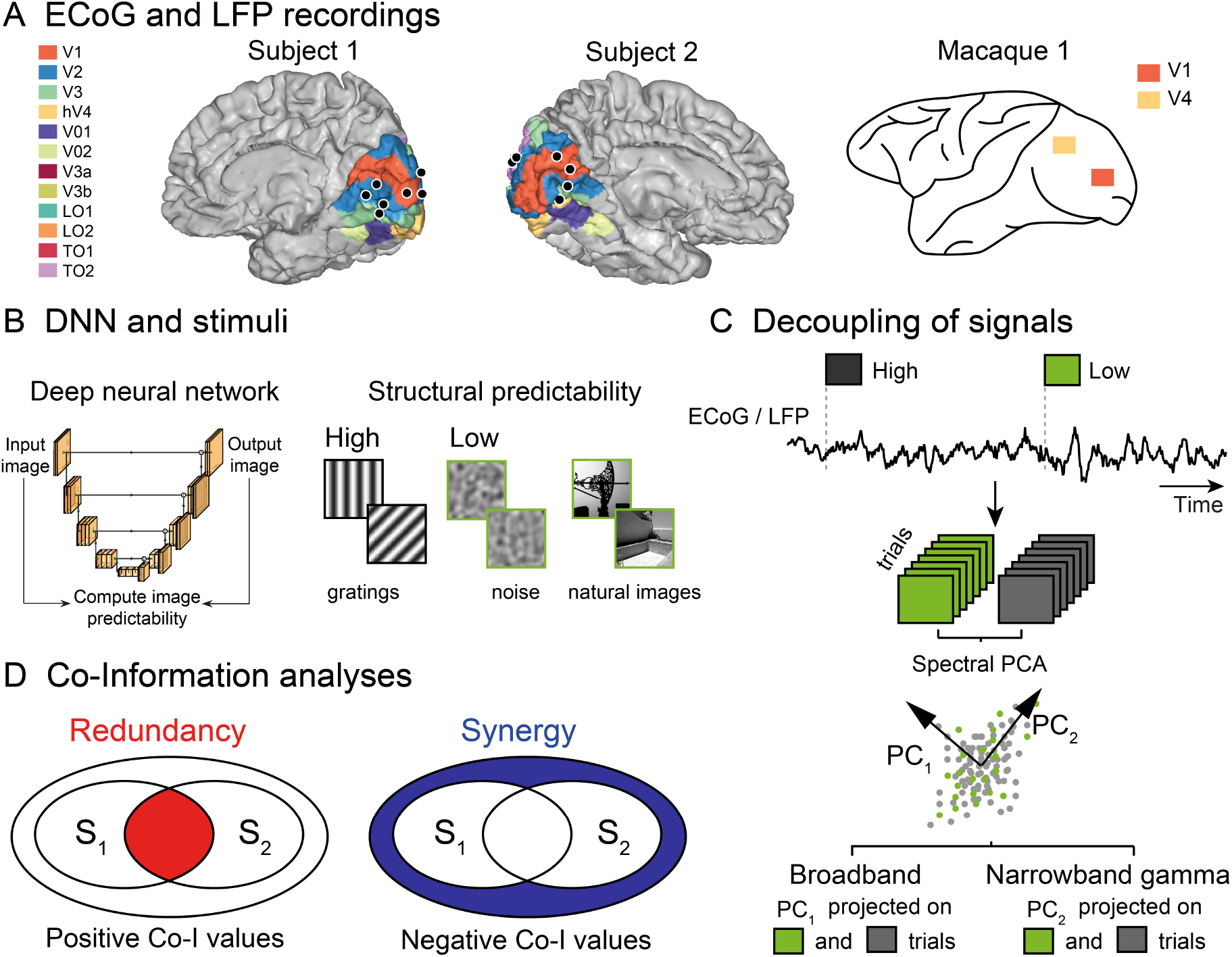
Redundancy and synergy analyses, stimuli, and decoupling neural ECoG and LFP responses. **(A)** Location of the electrodes implanted in Subject 1 (S1) and Subject 2 (S2) (black dots) rendered on estimates of early visual areas, and schematic representation of Macaque 1 (M1) brain with electrode locations (orange and yellow squares). **(B)** The spatial predictability of images was quantified using a technique based on self-supervised, deep neural networks (DNN; see Methods). In brief, a DNN is trained to predict visual images into the receptive fields (RFs). A mask of approximately the same size as the recording site’s RF is applied to an image. The image with the mask is then entered as an input to a DNN with a U-net architecture. This DNN predicts the full image, i.e., the image content behind the mask is filled in. Stimulus predictability is computed by comparing the ground-truth input image and the predicted image and then using it for network optimization during the training stage. The images were identical to those used in a prior ECoG study and grouped into several stimulus categories (S1 and S2: space, orientation, contrast, sparsity, and coherence; M1: orientation, and natural images). The images with the highest predictability score (high spatial homogeneity: gratings) were compared against those with the lowest predictability scores (low spatial homogeneity: noise and natural images). **(C)** Decoupling neural signals from ECoG and LFP data. Epochs corresponding to high homogeneity (black) and low homogeneity (green) images are used to compute the broadband (BB) response and narrowband gamma (NBG) response. BB signals are computed by reconstructing the time series of each stimulus category trial with the first spectral principal component (SPCA) of the ECoG/LFP signal; NBG are computed by reconstructing the same time series with the second principal component (SPCA) of the ECoG/LFP signal (see Methods). **(D)** Schematic representation of redundancy and synergy analyses computed using co-information. Each inner oval (S1 and S2) represents the mutual information between the corresponding ECoG/LFP signals and the stimuli category. The overlap between S1 and S2 represents the redundant information about the stimuli (red; left panel). The outer circle around S1 and S2 represents the synergistic information about the stimuli (blue; right panel). S1 and S2 can be signals from the same area (e.g. V1) or between areas (e.g. V1 and V2).

We focused on a feature of images that is known to be reliably encoded by NGB in both humans and macaques, namely the structural predictability of images (Hermes et al., 2019; Shirhatti et al., 2022; Uran et al., 2022; Vinck and Bosman, 2016). We operationalize structural predictability as the extent to which the image in the surround predicts the precise structure of receptive field inputs in the center Uran et al. (2022), which relates to the predictability of structural features such as orientations across space (Coen-Cagli et al., 2012; Hermes et al., 2019). Uran et al. (2022) have shown that, among many image features, structural predictability (as defined above) is the best predictor of NGB. This approach allowed us to compare the nature of information representations between BB and NBG in terms of synergy and redundancy. In both humans, we analyzed a previously reported dataset in which bandpass-filtered grayscale images were analyzed that varied in several parametric dimensions. We compared gratings to a set of images with low spatial predictability. By using a deep neural network (DNN) that predicts the visual input in the center from the surround (similar to (Uran et al., 2022), we verified that gratings had higher structural predictability than noise images (S1: p < 0.001, S2: p = 0.002; see Methods). In macaque M1, we analyzed gratings and grayscale natural images previously reported by Uran et al. (2022). We compared gratings to a set of natural images that had low structural predictability, as identified with the DNN (difference gratings and natural images, p < 0.001; see Methods).

### Decoupling broadband and oscillatory responses from the ECoG and LFP signals

To separate the oscillatory NBG from the non-oscillatory BB neural dynamics in the ECoG and LFP signals, we performed the spectral decoupling technique (Miller et al., 2009a; Miller, 2019) (Figure 1C). In brief, for each electrode in each subject (S1, S2) and macaque (M1), we computed spectral principal component analysis (SPCA) across the power spectra from all trials. The time series for each trial was then reconstructed using the weights of the first and second SPC, separately (see Methods; Figure S1 and Figure S2). In each electrode and subject (S1: 7 electrodes; S2: 6 electrodes; M1: 126 electrodes: 63 in V1 and 63 in V4; Figure 1A), we observed the characteristic BB first principal component reported in previous ECoG studies during visual and auditory processing (Canales-Johnson et al., 2021; Gelens et al., 2024; Miller, 2019; Miller et al., 2017; Sabra et al., 2019). The BB component spanned frequencies from 1 to 200 Hz without a characteristic peak (Figure 2A in dark green; Figure S1 and Figure S2).

**Figure 2:**
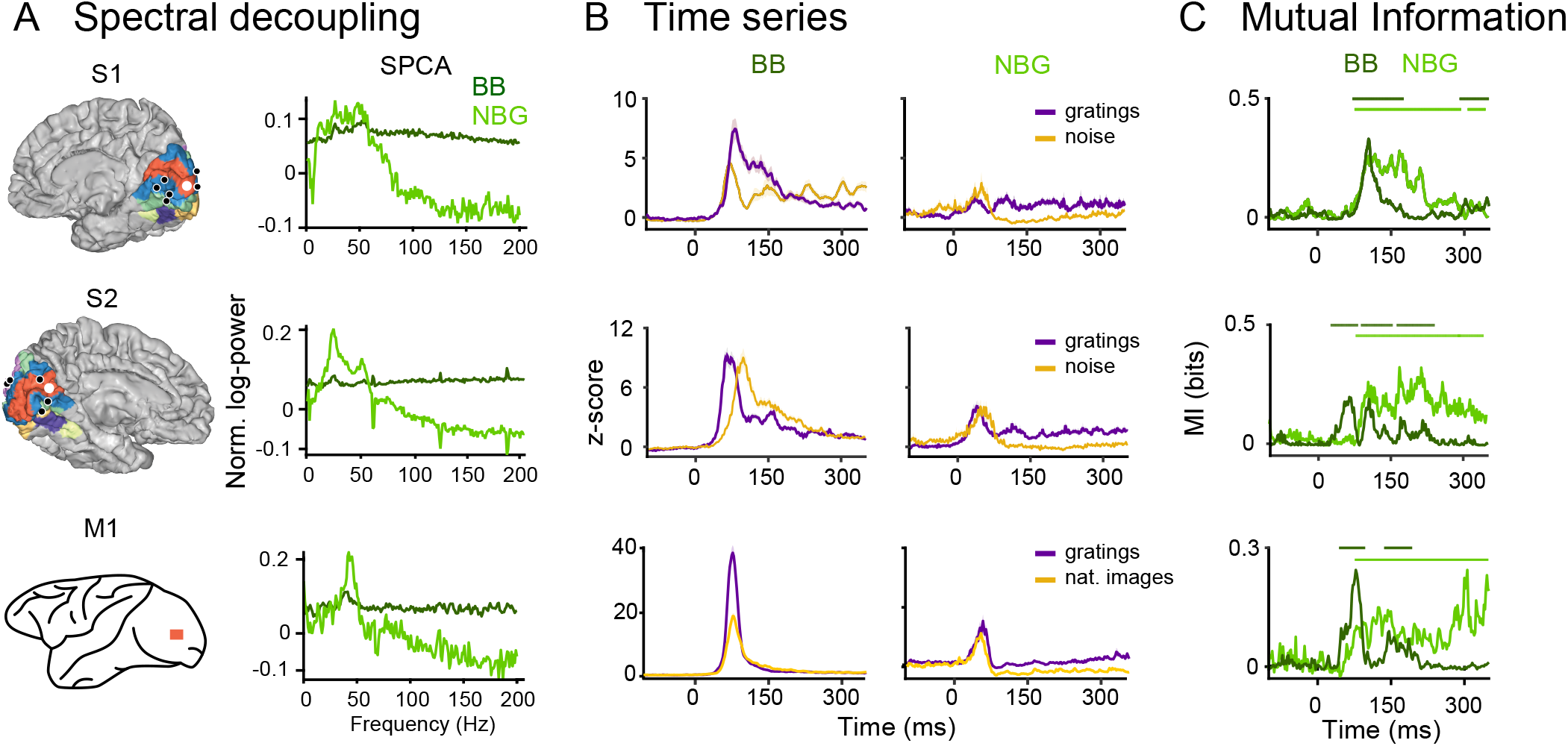
Spectral decoupling and mutual information analyses. **(A)** Spectral decoupling from a V1 electrode in Subject 1 (S1) and Subject 2 (S2) (in white), and from a V1 electrode in Macaque 1 (M1) (in orange). The magnitude (i.e., the normalized log-power) of the first (dark green) and second (light green) spectral principal component (SPCA) is shown between 1 and 200 Hz. The first SPCA is the non-oscillatory, broadband (BB) component, and the second SPCA is the oscillatory, narrow-band gamma (NBG) component. **(B)** Time-series average across gratings (purple), and noise (yellow) trials, reconstructed with the BB (left panel) and NBG components (right panel) for S1, S2, and M1. **(C)** Mutual information analyses between gratings (dark green) and noise images (light green) for S1 and S2, and between gratings and natural images for M1. Mutual information (in bits) quantifies the effect size of the difference between the corresponding image categories, representing the amount of information encoded in the signals about the stimulus difference. The significant time points after a permutation test are shown as bars over the MI plots.

In contrast, the second spectral component exhibited a narrowband profile, spanning frequencies in the range of the oscillatory, narrowband gamma band (NBG: ∼ 30-80 Hz), and it was observed in 4 out of 7 electrodes of S1, in 5 out of 6 electrodes of S2, and in 63 out of 63 electrodes in M1 (Figure 2A in light green; Figure S1 and Figure S2). While the BB response is induced by all images, including spatially heterogeneous images (Hermes et al., 2015a, 2019), the NBG response is predominantly induced by spatially homogeneous images such as gratings (Hermes et al., 2015a, 2019). Note that these induced BB and NBG changes are best appreciated after ∼100 ms when transient evoked responses have less influence on the signal. A representative electrode (area V1) for S1, S2, and M1 is depicted in Figure 2B for gratings (in purple), noise, and natural images (in yellow), reconstructed with the BB (dark green) and NBG components (light green). The results for all electrodes and datasets are depicted in Figure S3B and Figure S4B, and Figure S5. The power spectral density (PSD) for the selected electrodes and stimuli categories is depicted in Figure S3A and Figure S4A).

### Information about image predictability is encoded in BB and NBG signals

After decoupling the non-oscillatory BB and the oscillatory NBG responses from the ECoG and LFP signals, we quantified the amount of information encoded in the signals about image predictability using information-theoretic measures. To this end, we computed Mutual Information (MI) between trials corresponding to images with high and low structural predictability (Figure 2C) using the Gaussian-Copula Mutual Information (GCMI) estimator (Ince et al., 2017). Within the framework of information theory, MI is a statistical measure that quantifies the strength of the dependence (linear or nonlinear) between two random variables. It can also be seen as the effect size, quantified in bits, for a statistical test of independence.

Specifically, we computed MI between the neural response R (BB or NBG) and the stimulus category S (gratings vs noise in humans; grating vs natural image in macaque). MI quantifies, in bits, the reduction in uncertainty about S given R and was estimated with the Gaussian Copula MI framework (see Methods). This provides a continuous measure of category discriminability and is evaluated at each time point per electrode with FWER-controlled permutation testing. Thus, we used MI separately for the BB and NBG responses to quantify differences between image categories on a standardized effect-size scale. We observed significant MI encoded in the BB (dark green) and NBG (light green) signals after comparing gratings versus noise in S1 and S2, and gratings versus natural images in M1 (Figure 2C, and and Figure S3B). Although both BB and NBG responses encoded significant information about the structural predictability of images, we did observe some differences in latencies between the two types of dynamics (Figure 2C, and Figure S3B). Across electrodes and subjects (Figure S3B), the earliest significant MI effects appeared in the BB signals, while the MI effects in the NBG signals emerged later. Specifically, in S1, MI effects in the BB signals emerged as early as 28 ms after stimulus onset compared to the 60 ms onset of the NBG signals (Figure S3B; upper panel). Similarly, in S2, BB effects began at 48 ms while NBG effects began at 80 ms (Figure S3B; middle panel). In M1, BB effects began at 50 ms and NBG effects at 66 ms (Figure S3B; lower panel).

### Temporal redundancy in BB and NBG signals

The finding that BB and NBG signals encode information about image predictability raises the question of whether the information encoded within visual regions is shared (redundant) or complementary (synergistic) across time. To answer this question, we separately computed synergistic and redundant information across time using co-information (co-I) for the BB and NBG responses. Co-I tests whether two neural responses carry the same information (redundant) or if the relationship between them carries extra (synergistic) information about the stimulus: positive values indicate redundancy; negative values indicate synergy (see Methods and Figure 1D). Co-I analyses revealed temporal clusters of synergistic information (in blue) and redundant information (in red) across time within BB and NBG signals in S1 and S2 when contrasting gratings versus noise images (Figure 3).

**Figure 3:**
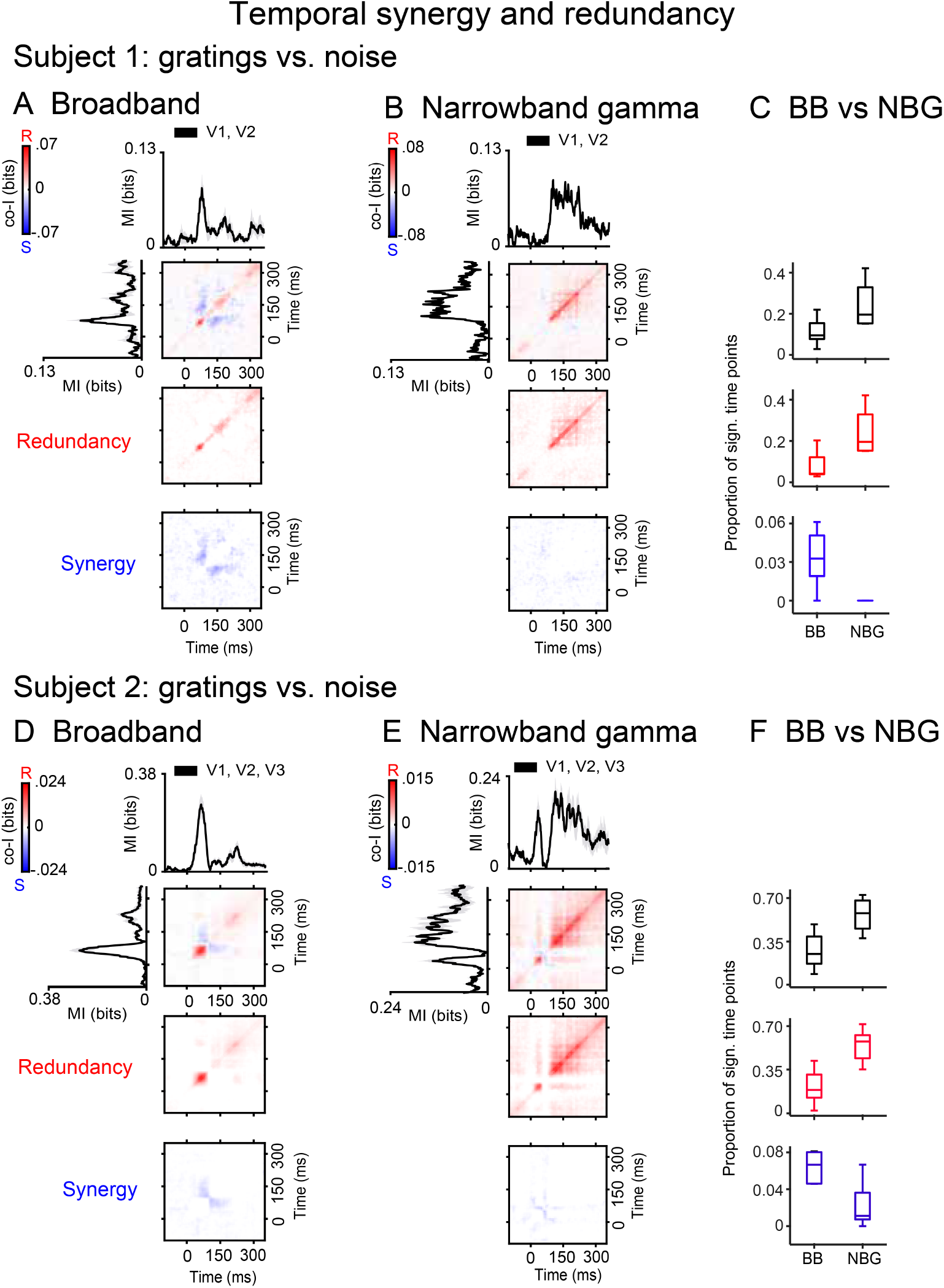
Temporal synergy and redundancy within visual areas in the human brain. Co-information revealed synergistic and redundant temporal patterns within the visual cortex in Subject 1 and Subject 2. (**A, D**) Co-information charts for BB signals. (**B, E**) Co-information charts for NBG signals. Black traces represent the MI between gratings and noise trials for the corresponding signals. Error bars represent the standard error of the mean (S.E.M) in V1 and V2 electrodes for S1 (**A, B**), and in V1, V2 and V3 electrodes for S2 (**D, E**). Temporal co-I was computed within visual areas across time points between -100 to 350 ms after image presentation. The average across individual recording sites is shown for the complete co-I chart (red and blue panel); for positive co-I values (redundancy only; red panel); and negative co-I values (synergy only; blue panel). (**C, F**) Box plots display the sample median alongside the interquartile range. Samples represent the percentage of significant time points observed in each electrode for co-I (black boxes), redundant (positive values; red boxes), and synergistic information (negative values; blue boxes) in the corresponding BB and NBG signals.

Regarding the dynamics of redundant information within BB and NBG signals, we observed different patterns of temporal redundancy for the two signals. Redundancy quantifies the degree to which two time points carry the same trial-by-trial information about the stimulus category (i.e., high vs. low structural predictability). This is conceptually similar to the temporal generalisation method (King and Dehaene, 2014), which quantifies the similarity of a multivariate predictive response pattern over time. Similarly, different temporal patterns of redundancy can reveal different signatures of information representation. In qualitative terms, a diagonal is a pattern described by King and Dehaene (2014) as a *chain*, where a sequence of discrete generators represent the stimulus over time. In contrast, a square pattern is a qualitative term that indicates a *sustained* response, which is more likely to result from a single process maintained over time.

In the case of redundancy, a diagonal pattern indicates a continually evolving representation of information, a representation that is distributed over time. In such a system, a downstream readout would benefit from integrating the signal over time, as later time points continue to provide new information. In the BB signals, diagonal redundancy was observed in S1 and S2 (Figure 3A,D) and M1 (Figure 5A).

Instead, a sustained block pattern was observed in the NBG signals. Redundancy over a large temporal region suggests a unitary information processing event in which the same information is obtained, whichever time point you sample from. In this case, a downstream processor would only need to observe one time point to get all the information, although integration might still have benefited from a signal-to-noise (SNR) ratio perspective. In the NBG signals, we observed sustained redundancy over time across the entire period after stimuli presentation in S1 and S2 (Figure 3B, E; and M1 (Figure 5B).

To directly compare potential differences in how BB and NBG signals encode redundant representations about image predictability, we quantified and statistically contrasted the temporal extension of redundancy between BB and NBG by a two-sided t-test on the percentage of significant time points per electrode (see Methods). We observed more temporal redundancy, as measured by the number of significant time points, in NBG compared to BB signals in S1 (t_(1,9)_ = 2.83; p = 0.019; n = 4 vs. n = 7; Figure 3C); and S2 (t_(1,12)_ = 3.90; p = 0.003; n = 5 vs. n = 6; Figure 3F); and M1: (t_(1,62)_ = 8.30; p < 0.001; n = 63 vs. n = 63; Figure 5C), and also more generally in co-I (S1: t_(1,9)_ = 2.30; p = 0.046; S2: t_(1,12)_ = 3.28; p = 0.009; M1: t_(1,62)_ = 7.70; p < 0.001).

### Temporal synergy in BB and NBG signals

In contrast to redundancy, temporal synergy within BB signals was observed off-diagonally between early and late time points in S1 and S2 (Figure 3A,D), and M1 (Figure 5B). This off-diagonal synergy has a characteristic pattern with two lobes originating from a common starting point aligned with each temporal axis (Gelens et al., 2024). As for redundancy, we can ask what system would lead to this profile. One possibility is that the neural system undergoes a discrete change of state at the time point corresponding to the origin of the two synergy nodes. This means that the signal obtained from any time point after the origin of the synergy nodes gives information about the state of the neural system, which improves the readout of the represented stimulus information. This is how synergy can arise between time points even when one does not carry any stimulus information directly (i.e., without significant MI at those time points).

To compare these synergistic patterns to the redundant ones, we quantified and statistically contrasted the temporal extension of synergy between BB and NBG signals. Again, we compared BB and NBG by a two-sided t-test on the percentage of significant time points per electrode (see Methods). Interestingly, and in contrast to temporal redundancy, we observed more temporal synergy in BB compared to NBG signals in S1 (t_(1,9)_ = 3.06, p = 0.017; n = 7 vs. n = 4; Figure 3C); and S2 (t_(1,12)_ = 3.27, p = 0.012; n = 6 vs. n = 5; Figure 3F), and M1 (t_(1,62)_ = 2.42, p = 0.016; n = 63 vs. n = 63; Figure 5C).

### Spatiotemporal redundancy in BB and NBG signals

The observation that BB and NBG signals convey distinct types of information about image predictability within visual regions raises the question of whether BB signals and NBG signals would primarily encode synergistic and redundant information *between* visual areas, respectively.

Regarding redundancy, we observed different spatiotemporal patterns between areas in the BB and NBG signals. Similarly to the results observed within areas, interareal interactions in the NBG signals were highly sustained over time compared to the redundant patterns observed between BB signals, which exhibited redundancy along the diagonal (Figure 4). In S1, redundant interactions were observed between areas V1-V2 (Figure 4A,B); and between areas V1-V2/V3 in S2 (Figure 4D,E), and between V1 and V4 in the case of M1 (Figure 6).

**Figure 4:**
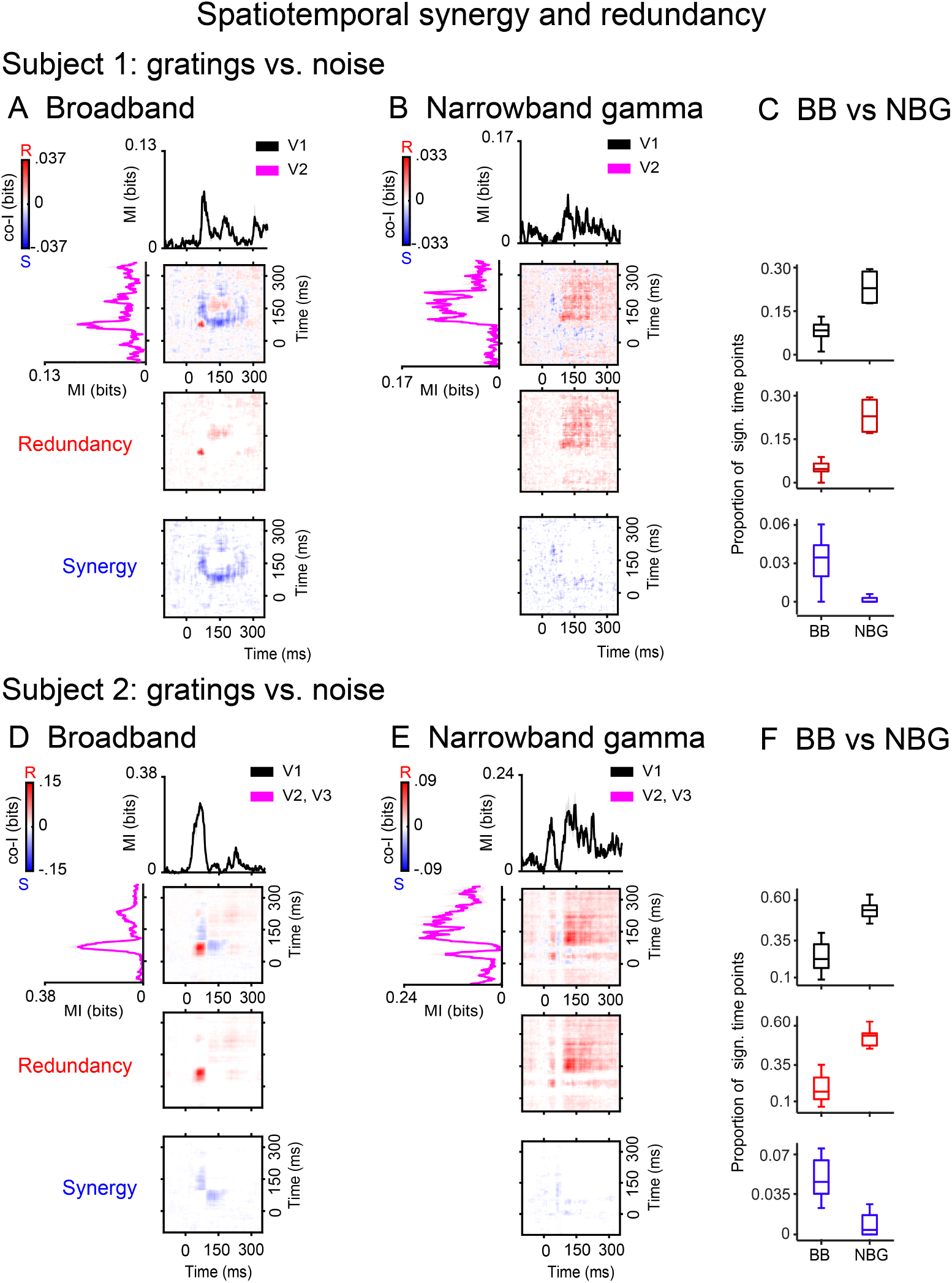
Spatiotemporal synergy and redundancy between visual areas in the human brain. Co-information revealed synergistic and redundant temporal patterns between visual areas in Subject 1 and Subject 2. (**A, D**) Co-information charts for BB signals. (**B, E**) Co-information charts for NBG signals. Black traces (electrodes in V1) and magenta traces (electrodes in V2, and V3) represent the MI between gratings and noise trials. Error bars represent the standard error of the mean (S.E.M) between the corresponding electrodes. Co-I was computed between each pair of electrodes (V1-V2 for S1; V1-V2 and V1-V3 for S2) and across time points between -100 to 350 ms after image presentation. The average across inter-areal electrode pairs is shown for the complete co-I chart (red and blue panel); for the positive co-I values (redundancy only; red panel); and the negative co-I values (synergy only; blue panel). (**C, F**) Box plots display the sample median alongside the interquartile range. Samples represent the percentage of significant time points observed in each electrode for co-I (black boxes), redundant (positive values; red boxes), and synergistic information (negative values; blue boxes) in the corresponding BB and NBG signals.

We quantified and statistically contrasted differences between BB and NBG signals as described above, this time between the corresponding visual areas. We compared BB and NBG by a two-sided t-test on the percentage of significant time points per electrode pair (see Methods). Again, we observed higher temporal redundancy in NBG compared to BB signals in S1 (t_(1,9)_ = 3.90; n = 6 pairs vs. n = 21 pairs; p = 0.003; Figure 4C); and S2 (t_(1,12)_ = 7.39; p < 0.001; n = 10 pairs vs. n = 15 pairs; Figure 4F), and also in co-I (S1: t_(1,9)_ = 6.46; p < 0.001; S2: t_(1,12)_ = 6.10; p < 0.001).

In the case of M1, since we did not observe NBG signals in V4, we only report V1-V4 interactions in the BB signals (Figure 6). To quantify whether spatiotemporal redundancy and co-I were statistically significant above chance, we performed permutation tests on the co-I and redundancy charts separately (see Methods). Significant redundancy between V1 and V4 was observed in 3597 out of 3969 electrode pairs (90.6%). Similarly, we observed significant co-I in 3686 out of 3969 V1-V4 electrode pairs (92.8%).

### Spatiotemporal synergy in BB and NBG signals

In contrast, and consistent with the findings within regions, interareal synergistic information was mainly observed in the BB signals in S1 and S2 (Figure 4A,D) and in M1 (Figure 6A). Across species, interareal synergy was observed off-diagonally between the peak of the MI signal in one region and later time points of the MI signals in another region (S1: Figure 4A; S2: Figure 4D; M1: Figure 6). As per redundancy, we contrasted the spatiotemporal synergy between BB and NBG signals between the corresponding visual areas. We observed higher spatiotemporal synergy in BB compared to NBG signals in S1 (t_(1,9)_ = 3.42; p = 0.004; Figure 4C); and S2 (t_(1,12)_ = 4.72; p < 0.001; Figure 4F).

In the case of M1, to quantify whether spatiotemporal synergy between V1 and V4 was significant above chance, we performed permutation tests in the synergy charts (Figure 6). Significant spatiotemporal synergy between V1 and V4 was observed in 1721 out of 3969 electrode pairs (43.4%). Note that co-I is non-directional; the apparent asymmetry reflects earlier V1 information and later V4 information, producing an off-diagonal synergistic pattern aligned with the feedforward delay. We also note an apparent difference in the temporal extent of inter-areal BB interactions across species. In humans (Figure 4), BB synergy and redundancy between V1-V2/V3 are relatively confined in time, whereas in macaque (Figure 6) synergy between BB responses and redundancy between V1 and V4 span a broader post-stimulus window. Two factors likely contribute. First, in macaque V4, we did not detect a reliable NBG component (0/63 electrodes; see Methods), so Figure 6 reflects only BB-BB interactions, whereas Figure 4 compares BB and NBG communication directly. Second, the macaque comparison (gratings vs natural images) produces relatively long-lasting MI in the BB response in area V4, and this increased MI in each area constrains spatiotemporal co-I. This may correspond to more temporally extended BB co-I without changing the qualitative conclusion that BB communication is primarily synergistic.

## DISCUSSION

In the visual system, BB fluctuations and NBG oscillations are commonly superimposed (Hermes et al., 2015a, 2019; Ray and Maunsell, 2011). We note that BB activity is conventionally defined empirically as the absence of spectral peaks in the PSD and a 1/f structure in the PSD (Donoghue et al., 2020; Miller et al., 2009a); we contrast BB with oscillatory activity, which is reflected by clear spectral peaks in the PSD caused by frequency-band-limited synchronous activity. Studies indicate that BB activity primarily reflects the contribution of aperiodic synaptic currents. These aperiodic currents contain contributions of both uncorrelated and correlated synaptic inputs (Einevoll et al., 2007; Lindén et al., 2010). At higher frequencies, spikes may also directly contribute energy to the LFP, particularly when the neurons are located near the electrode (Anastassiou et al., 2015; Buzsáki et al., 2012). The energy of BB activity is expected to correlate with firing rates, which is consistent with empirical reports (Ray and Maunsell, 2011). Importantly, BB and narrowband oscillatory activity overlap in frequency space and thus require decomposition techniques to be disentangled. This is possible because of their different underlying generative processes and because they show differential correlations with various stimulus factors (Hermes et al., 2015a, 2019; Jia et al., 2013b; Peter et al., 2019; Ray and Maunsell, 2011; Uran et al., 2022)

Despite their different neural generators, it remains an open question what role BB and NBG dynamics play in communication and whether they play distinct and complementary roles in encoding and transmitting information (Vinck et al., 2025, 2023). We investigated whether broadband and oscillatory dynamics play different roles in encoding and transmitting synergistic and redundant information. The redundancy metric captures the shared (or common) information that a set of signals encodes about a stimulus variable. On the other hand, the metric of synergy quantifies the complementary (or extra) information that a set of signals encodes as a whole, above and beyond the information encoded by the sum of the isolated signals (Gelens et al., 2024; Ince et al., 2017).

To test this hypothesis, we recorded from multiple visual areas in the macaque and human cortex and decomposed signals into broadband and narrowband gamma activity. Broadband signals already carried substantial information about visual images at short latencies after stimulus onset, while narrowband activity carried information in a sustained manner, consistent with previous hypotheses (Vinck et al., 2023). We show broadband signals encoded significant synergistic information about visual images, whereas narrowband signals were predominantly redundant.

### Redundant versus synergistic interactions

Empirical studies in several species and sensory modalities have shown that synergy and redundancy have functional relevance for encoding stimuli and task variables (Blume et al., 2026; Combrisson et al., 2024; Delis et al., 2022; Francis et al., 2022; Gelens et al., 2024; Greco et al., 2024; Koçillari et al., 2023; Nigam et al., 2019; Olivares et al., 2025; Park et al., 2018; Varley et al., 2023). In the auditory system, broadband signals between the primary auditory cortex (A1) and frontal areas exhibit synergistic encoding of prediction errors (Gelens et al., 2024). Computational modeling suggests that such synergistic relations between distributed prediction error signals can result from nonlinear recurrent dynamics between regions (Gelens et al., 2024). In the visual system, laminar recordings in the primary visual cortex (V1) indicate that synergistic interactions can decode visual stimuli more effectively than redundant interactions, even when faced with noise and overlapping receptive fields (Nigam et al., 2019). In contrast, LFP recordings across the olfactory system show that narrowband oscillations exhibit higher levels of redundant information about odorant stimuli (Olivares et al., 2025). In human MEG recordings, theta oscillations (3-7 Hz) in the temporal cortex represent auditory and visual input redundantly during the multimodal integration of audio-visual speech signals (Park et al., 2018). Contrarily, the same narrowband oscillations in the motor and inferior temporal cortex represent the inputs synergistically (Park et al., 2018). What mechanisms account for the differences in synergy and redundancy observed here? We posit that synergy results from nonlinear recurrent interactions between nodes, such that information resides in the joint pattern of activity across nodes. In support of this idea, a recent study shows that adding recurrent interactions between nodes causes a switch from redundant to synergistic representations (Gelens et al., 2024). However, we would like to emphasize that synergy in our study is a statistical descriptor of joint neural interactions, not a mechanistic model of the underlying neurophysiology. Mechanistic inferences, therefore, require converging evidence from modelling and/or targeted causal perturbations. In support of our interpretation of synergy as recurrent processing between brain areas, Gelens et al. (2024) showed that in a brain-constrained neurocomputational model matched to marmoset ECoG, temporofrontal synergy emerged only when strong long-range recurrent (feedback and feedforward) *jumping* connections were included; purely feedforward or locally recurrent variants did not produce synergy. Thus, this observed inter-areal synergy is consistent with an interpretation of recurrent processing across cortical levels. We note, however, that synergy can also arise from global state fluctuations (e.g., attention/arousal) or other non-stimulus-specific factors, so we present this as a candidate mechanism to be tested with targeted perturbations and further modelling.

By contrast, redundancy in narrowband oscillatory signals may simply reflect that linear transformations yield communication at the same frequency, and only relay information. Thus, linear information transmission leads to redundancy and signal transfer within the same frequency band. We have previously shown that a sending and receiving neural population will naturally exhibit coherent activity because spiking activity in a sending area causes synaptic potentials in the same area and highly correlated synaptic potentials in another receiving area (at a delay) (Dowdall et al., 2023; Dowdall and Vinck, 2023; Schneider et al., 2021). Hence, coherence can be a consequence of linear signal transmission rather than the cause of inter-areal communication. Furthermore, we showed that narrowband gamma-band signals may not impact spiking activity in downstream areas, such that they primarily contribute to inter-areal coherence via the generation of subthreshold synaptic potentials in downstream neurons (Schneider et al., 2023; Spyropoulos et al., 2024). Finally, gamma oscillations in the visual cortex are well approximated by linear models (damped harmonic oscillators) (Spyropoulos et al., 2022), such that integration within the gamma-frequency band may take place via linear resonance rather than nonlinear entrainment mechanisms (Lewis et al., 2021). Thus, the redundancy between narrowband signals shown here may reflect linear transmission between sites. Even if there are nonlinear interactions within the same frequency band, communication within the same frequency band may typically involve relaying information, leading to redundancy rather than extracting patterns via nonlinear transformations. In fact, modeling studies investigating selective communication via coherence (Communication-through-Coherence; CTC) have focused on scenarios where information is relayed between areas (Akam and Kullmann, 2012).

### Functional implications

Our findings suggest that broadband and narrowband dynamics have complementary functions in the encoding and transmission of information. We posit that the characteristics of broadband activity may be ideally suited for transmitting synergistically encoded information. Notably, synergistic information represented by two populations *X* and *Y* typically requires nonlinear read-out mechanisms in a receiving area *Z*. For example, in the case of the classic example of a synergistic function, the XOR-gate (Down and Mediano, 2025; Timme et al., 2014), a receiver should compute a function *g*(*X, Y*) to extract the information encoded in the joint activity pattern rather than an additive function *f* (*X*) + *h*(*Y*). In other words, a synergistic code in *X* and *Y* might imply that *X* and *Y* exert non-separable, nonlinear influences. Nonlinear integration mechanisms in the cortex may predominantly rely on broadband signals that allow for interactions across frequencies (Vinck et al., 2023), which agrees with our finding that broadband activity predominantly carries synergistic information. Notably, this broadband activity carried information especially early after stimulus onset, while oscillatory activity carried information substantially later. We speculate that early, largely transient BB synergy might reflect rapid perceptual inference supported by nonlinear, recurrent interactions across neural populations (Vinck et al., 2025). In our results, BB synergy typically emerges between early and later time points off-diagonally, consistent with the idea that a brief input-driven event pushes the system into a new multi-areal attractor state, in which information about the stimulus is encoded in the joint activity pattern across nodes rather than at any single site. In such a regime, readout requires nonlinear integration of population activity (e.g., XOR-like computations), which is naturally captured by synergistic information.

Visual processing is a highly nonlinear process (Cohen et al., 2020; DiCarlo et al., 2012) that depends partially on recurrent interactions (Kar et al., 2019). In fact, clusters of pixels collectively provide synergistic information about an object’s identity. This is because individual pixel values alone offer very little insight into what the object is. However, when these pixel values are considered together, the relational information in the pattern improves object recognition. Therefore, a system that performs perceptual inference by nonlinearly integrating information over space can be expected to exhibit synergistic representations. This principle can even be seen when nodes are explicitly initialized in an oscillatory manner: A recent modeling study has shown that a recurrent neural network that is initialized with oscillating nodes gradually desynchronizes and increases its heterogeneity across nodes during the learning of a pattern recognition task, indicating that cross-frequency relations are required for performing a complex pattern recognition task (Effenberger et al., 2025).

Interestingly, early theories on perceptual integration emphasized that perceptual binding relies on synchronized oscillatory firing (Singer, 1999). In such a case, the information about the object (i.e., representing the same object) would be shared across nodes, leading to redundant representations. An alternative possibility is that perceptual binding relies on synergistic representations that emerge from nonlinear interactions between nodes. Indeed, previous work suggests that nonlinear interactions across distributed nodes rather than linear interactions or spectral power distinguish different perceptual states, given a constant sensory input (Canales-Johnson et al., 2023, 2020). Here, an interesting connection can be made with theories of consciousness emphasizing that consciousness reflects integrated information (Tononi et al., 2016), as synergistic representations among nodes (e.g. *X* and *Y*) imply intrinsic causal influences that are nonseparable to extract the information.

Compared to broadband information, which was particularly strong at earlier phases and became weak during late stimulus phases, information at later stimulus phases was carried predominantly by narrowband gamma activity, with information represented redundantly. This may suggest that information is sustained with a higher degree of redundancy across space and time after the initial stimulus phase in which information is synergistically encoded via broadband activity. Indeed, narrowband oscillatory activity may be functionally well suited to sustain representations over time in an efficient and stable manner (Chalk et al., 2016; Vinck et al., 2025). Notably, the emergence of narrowband gamma activity may be restricted to a subset of visual stimuli, while broadband activity may be ubiquitous across stimuli. In particular, narrowband gamma activity emerges predominantly, or exclusively, for a subset of natural stimuli (Hermes et al., 2015a,b, 2019; Peter et al., 2019; Uran et al., 2022). Moreover, gamma is generally strongest for low-dimensional visual stimuli with a high degree of predictability in visual inputs across space and time (Hermes et al., 2019; Peter et al., 2019; Shirhatti et al., 2022; Uran et al., 2022; Vinck and Bosman, 2016). Hence, narrowband gamma may emerge primarily for stimuli with a high degree of spatial redundancy, which aligns with the present finding that narrowband gamma carries redundant information representations.

It is possible that differences in neural activity between stimulus categories may relate to microsaccadic activity. We analyzed microsaccades in the available eye-tracking dataset (macaque M1) across different image categories. These results do not reveal an association with microsaccades (Figure S7). Regardless, in this study, we do not make a claim regarding the neural mechanisms mediating the effect of structural predictability on neural activity. To our knowledge, the mechanisms underlying such contextual modulation are not fully established. For example, there are several biophysical models (Jadi and Sejnowski, 2014; Tahvili et al., 2025), and indications that SOM neurons are more active for unpredicted stimuli (Keller et al., 2020), which in turn promote gamma oscillations. However, direct evidence for these mechanisms in primates and humans is lacking, and differences between rodents and primates warrant consideration (Onorato et al., 2020). Microsaccades are an unlikely mechanism for the relationship between structural predictability and NBG, for various reasons: Classic findings on gamma oscillations with size tuning have been made under anesthesia in monkeys (Jia et al., 2013a,b), where one can rule out eye-movement-related changes in activity, and the difference in gamma between noisy and grating images is also found in anesthesized, paralysed cats(Zhou et al., 2008). In Figure 1 and Figure Supplement 3 of Peter et al. (2019), we have shown that V1 gamma is not affected by restricting the analyses to periods outside microsaccades. Hence, the literature suggests that differences in NBG between structurally predicted and unpredicted stimuli are not driven by microsaccades. Likewise, gamma responses in monkey visual cortex are transiently disrupted by saccades during free viewing but then emerge image-specifically after each saccade (Brunet et al., 2014).

Finally, one may argue that the BB component, by spanning a wide frequency range, necessarily contains more information or synergy. This is unlikely for two reasons. First, synergy depends on excess joint information beyond the sum of individual MIs. It therefore cannot be produced by a higher SNR or wider spectra alone. Second, our pipeline reduces both BB and NBG to matched one-dimensional time series before the MI and co-I computations. Hence, the information measures capture trial-by-trial complementarity in the time domain rather than the spectral domain. Consistent with this, we observed BB off-diagonal synergy within and between areas. In contrast, NBG exhibited sustained redundancy even when its MI was strong, indicating distinct computational roles rather than purely bandwidth differences.

To sum up, the present work shows a new principle: synergistic information in broadband activity and redundant information in gamma-band oscillations. This highlights a potential dual role for these two features of cortical dynamics in sensory processing, with broadband dynamics subserving nonlinear pattern recognition and oscillations subserving information sharing, contributing to robustness.

## Authorship contributions

Conceptualization: AC-J and DH. Data analysis: LR, CU, FB, JÄ, and AC-J. Human recordings: DH, KJM, MAJ. Macaque data: CU, MV. Software and methods for human and macaque data: AC-J, CU, MV, RI, and DH. Visualization: LR, CU, JÄ, and AC-J. Writing and editing: AC-J, LR, MAJ, JÄ, CU, MV, RI, and DH. Supervision: AC-J.

## Acknowledgments

ACJ is funded by a Swedish Research Council Project grant (VR; 2025-03245), an ANID/FONDECYT Regular (1240899) and ANID/FONDECYT Regular (1251273) research grants. DH is funded by the National Eye Institute of the National Institutes of Health under Award Number R01EY035533 and R01EY023384. The content is solely the responsibility of the authors and does not necessarily represent the official views of the National Institutes of Health. This project was funded by the Neurological Foundation of New Zealand Grant (1839RF, to Corinne Bareham) and the Royal Society of New Zealand Marsden (MAU2011, to Dr. Corinne Bareham). MV and CU were financed by an ERC starting grant (850861) SPATEMP, DFG VI Grants (908/5-1 and 908/7-1), an NWO VIDI Grant, and the Dutch Brain Interface Initiative (DBI2).

## Data availability

The original human data used in this study is publicly available at: https://osf.io/q4yad.

## Code availability

The MATLAB/Python toolbox GCMI (Gaussian Copula Mutual Information) used in this study is publicly available at: https://github.com/robince/gcmi.

## METHODS

### Data acquisition

#### Human data

The human data analyzed in this study consisted of ECoG recordings in two human subjects. The data acquisition details are reported in full in Hermes et al. (2019), and data is available here: https://osf.io/q4yad. Subjects gave informed consent and the study was approved by the Stanford University IRB and the ethics committee at the University Medical Center Utrecht in accordance with the 2013 provisions of the Declaration of Helsinki.

This study modelled the broadband and gamma responses in early visual areas to 86 different stimuli varying across spatial location, orientation, contrast, sparsity, and coherence dimensions. ECoG potentials were recorded at 1528 Hz in the left hemisphere of S1 and the right hemisphere of S2 using 128-channel Tucker Davis Technologies and Micromed Ephys systems and resampled to 1000 Hz for computational purposes. The localization of electrodes was achieved through the co-registration of pre-operative MRI with a computer tomography (CT) scan, which was performed after the electrodes had been implanted. In each human subject, some electrodes showed overlapping population receptive fields. For each electrode, the location (x,y) and size (sigma) of the population receptive field (pRF) were taken from the previous calculation of a second-order contrast (SOC) model (Hermes et al., 2019), representing the location of the image to which the electrode responds. The pRFs of each electrode used in S1 and S2 can be found in Figure S6.

#### Macaque data

We also analyzed LFP data from one macaque (M1) previously reported by Peter et al. (2019) and Uran et al. (2022). M1 was positioned 83 cm in front of a 22-inch 120 Hz LCD monitor. The animal was implanted with a Utah array with 64 microelectrodes (inter-electrode distance 400 µm, half of them with a length of 1 mm and half with a length of 0.6 mm, Blackrock Microsystems), and a reference wire was inserted under the dura towards the parietal cortex.

The monkey self-initiated trials by fixating on a small fixation spot presented at the center of the screen and had to maintain fixation during the entire trial. Trials were aborted during which the eye position deviated from the fixation spot by more than 0.8 visual deg radius. Correct trials were rewarded with diluted fruit juice. We analyzed only correctly performed trials. Recordings were made using a Tucker Davis Technologies (TDT) system. The data were filtered between 0.35 and 7500 Hz (3 dB filter cutoffs) and digitized at 24414.0625 Hz (TDT PZ2 preamplifier). Line noise was removed using a two-pass 4^th^ order Butterworth bandstop filters between 49.9-50.1, 99.7-100.3, and 149.5-150.5Hz. We downsampled the LFPs to ≈ 1.02kHz using Matlab’s decimate.m function with the default settings, resulting in a 3dB low-pass cut-off frequency of 406.90 Hz. All procedures complied with the German and European regulations for the protection of animals and were approved by the regional authority. For further details see (Peter et al., 2019; Uran et al., 2022). Note that M1 is referred to as Monkey H in these studies.

### Stimuli and task

Human subjects were repeatedly presented with each type of stimulus image (15 trials per image), presented in a randomised order for a period of 500 ms, presented at an approximate visual angle of 20 degrees of visual angle left-to-right, with a baseline period between each presentation of the same duration, while maintaining fixation on a dot in the center of the screen. During the ECoG experiments for S1 and S2, images were displayed on a computer screen placed in front of the participant. Images were presented for 500 ms (stimulus period), followed by a 500 ms gray screen (baseline period). A small fixation dot was present at the center of the screen throughout the entire experiment. Stimuli consisted of 86 gray-scale patterns with gratings, curved lines, and noise patterns with 3 cycles per degree spatial frequencies. These patterns varied across well-controlled image dimensions, including spatial aperture, orientation, contrast, sparsity, and coherence (Hermes et al., 2019).

### Image predictability analysis

For both human and macaque subjects, we used an object-recognition convolutional neural network (OR-CNN) developed by Uran et al. (2022) to quantify the spatial predictability of each image channel (i.e., the extent to which the content of an area can be predicted from its spatial surroundings). Images were first cropped from their original size of 880 × 880 px to 224 px, centered on each electrode’s pRF. To calculate predictability scores per electrode and image type, we adapted the pred.py script from the model’s Unet-Pred repository, using the same weights and hyperparameters. Each image was masked 26 pixels from its center in width and height (radius) and fed to the UNet model with one step per image.

LPIPS (Learned Perceptual Image Patch Similarity) predictability scores were then calculated for each image by comparing the model’s predicted value for the central area with the true pixel values within the region. LPIPS predictability scores measure how different the predicted center of an image is from its true center, based on the surrounding context. Low scores indicate that the prediction is similar to the true center. High scores mean the prediction is dissimilar to the true center. Hence, according to our definition of image predictability, low scores indicate high structural predictability, while high scores indicate low structural predictability.

Thus, to identify images with the highest and lowest structural predictability in human subjects S1 and S2, we first ranked all images by their predictability scores. Oriented grating images (80 images) showed the lowest scores (i.e., high structural predictability). Noise images (80 images) showed the highest scores (i.e., low structural predictability). We then selected these two image categories for further analysis. As a result, the MI and co-I analyses were performed on 80 gratings and 80 noise trials per subject. To confirm that the predictability scores resulted in significant differences between categories, we used a two-sided t-test to compare the scores of the pre-selected images. Gratings showed the highest image predictability (i.e., lowest scores) in S1 (mean = 0.171; SD = 0.081) and S2 (mean = 0.179; SD = 0.082), and noise showed the lowest predictability (i.e., highest scores) in S1 (mean = 0.343; SD = 0.304) and S2 (mean = 0.307; SD = 0.274) (Figure 1B). As expected, we observed higher predictability for gratings compared to noise images across images and electrodes in S1 (t_(1,110)_ = 4.101, p < 0.001) and in S2 (t_(1,94)_ = 3.08, p = 0.002).

Unlike the human participants, in the original study by Uran et al. (2022), the macaque M1 was shown a set of natural images across 20 sessions. A total of 340 natural images were presented (20 trials per image). In each of the first four recording sessions, M1 was also shown a single oriented grating (20 trials per session). To identify images with the highest and lowest structural predictability in M1, we used LPIPS to determine the highest and lowest predictable natural stimuli in each session. This resulted in 80 gratings (20 images x 4 sessions = 80 images) and 80 natural-image trails (4 images with the lowest scores per session x 20 sessions = 80 images) being selected for further MI and co-I analyses. When comparing the predictability scores, we observed higher predictability for gratings (mean = 0.009; SD = 0.013) than for natural images (mean = 0.244; SD = 0.146) across images and electrodes (t_(1,470)_ = 24.45, p < 0.001).

The OR-CNN/LPIPS pipeline was used solely to rank stimuli by predictability (LPIPS) within each dataset; no DNN features are used in neural analyses. We used the published UNet (Uran et al., 2022) with fixed weights/hyperparameters and centered inputs on each electrode’s pRF. The neural MI/co-I effects reported here replicate across distinct image ensembles (band-passed gratings/noise in humans; gratings vs natural images in macaque), indicating that our conclusions do not hinge on model-specific training priors.

### Separating BB from NBG signals

Many standard methods that extract spectral power in predefined frequency bands cannot reliably dissociate broadband (BB) activity from narrowband oscillations (e.g., NBG). For instance, estimating gamma-band power in the 30-80 Hz range with a bandpass filter will generally mix two distinct contributions: (i) genuine NBG oscillations and (ii) the portion of the broadband component that falls within 30-80 Hz (Donoghue et al., 2020; Miller et al., 2009a) Moreover, several recently proposed approaches are optimized for power spectra that are averaged across trials (Hermes et al., 2015a, 2019) or computed over relatively long time windows (Donoghue et al., 2020). These methods are therefore less suited when fine-grained temporal precision is critical for computing information representations, as in the present study. In contrast, the Spectral Decoupling method (Canales-Johnson et al., 2021; Gelens et al., 2024; Miller et al., 2009a; Miller, 2019; Sabra et al., 2019) based on spectral principal component analysis (SPCA) approach operates at the level of trial- and time-resolved spectra and exploits covariance across frequencies to identify components that capture independent fluctuations in broadband and narrowband activity. This makes SPCA particularly well-suited to disentangle BB and NBG contributions with the temporal precision required for our analyses.

For each electrode, we computed a power spectral density (PSD) for every trial and stacked these spectra into a frequency-by-trials matrix *P*(*f, q*), where *f* indexes frequency bins (retaining positive frequencies up to 200 Hz) and *q* indexes trials (*q* = 1, …, *N*_*q*_). We normalized each frequency bin by its mean power across trials and applied a natural-log transform:

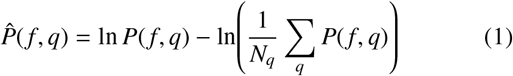

We then formed a frequency-frequency covariance matrix across trials,

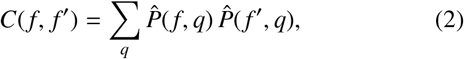

and performed eigendecomposition,

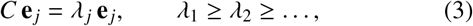

yielding orthogonal spectral principal components (SPCs) **e** _*j*_(*f*) ordered by explained variance. In our datasets, the first component (SPC1) captured broadband (aperiodic) modulations spanning 1–200 Hz, while the second component (SPC2) isolated a gamma-band spectral motif concentrated in the 30-80 Hz range.

To obtain a component-specific univariate time series for each trial, we computed a time-frequency representation using a bank of complex Morlet wavelets (5 cycles per frequency). Let *V*(*f, t, q*) denote the complex wavelet coefficient at frequency *f*, time *t*, and trial *q*. We computed band-power over time, discarded convolution edge samples, normalized each band to unit mean over time, and applied a natural-log transform:

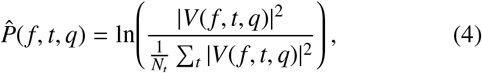

where *N*_*t*_ is the number of time samples in the analyzed window. We then projected the log-power matrix onto each SPC vector to yield a single component time course:

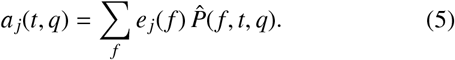

We treated *a*_1_(*t, q*) as the broadband (BB) component time course and *a*_2_(*t, q*) as the narrowband gamma (NBG) component time course. These component time series were epoched from −100 ms to 350 ms around stimulus onset and baseline-corrected by subtracting the mean of the pre-stimulus interval (−100 to 0 ms). For macaque data, reconstructed time courses were downsampled to 500 Hz to match the human sampling rate used in downstream analyses.

We identified NBG components using a run-length threshold applied independently to each spectral principal component (SPC) and electrode. For each SPC, we evaluated the frequency range 30-80 Hz (51 samples). An NBG component in a given electrode was marked present if this window contained a consecutive run spanning at least 75% of the window with magnitudes strictly greater than 0 (i.e., ≥ 39 consecutive samples within 30-80 Hz). Applying this criterion independently yielded a single NBG component in 4/7 electrodes in S1, 5/6 electrodes in S2, 63/63 electrodes in M1 (V1), and 0/63 electrodes in M1 (V4). Across S1, S2, and M1, the NBG component corresponded to SPC2 in all cases.

Similarly, the BB components were selected with the same procedure over 1-200 Hz (200 samples). A BB component in a given electrode was marked present if this window contained a consecutive run spanning at least 75% of the window with magnitudes strictly greater than 0 (i.e., ≥ 150 consecutive samples within 1-200 Hz). Using this criterion, we found a single BB component in 7/7 electrodes in S1, 6/6 electrodes in S2, and 63/63 electrodes in M1 in both V1 and V4. The BB component corresponded to SPC1 across S1, S2, and M1. The average across channels for NBG and BB components following this procedure is depicted in Figure S1. The projection of SPC1 and SPC2 on the time-frequency axis for gratings stimuli for S1, S2, and M1 is shown in S2.

### Mutual Information analyses

Within the framework of information theory, Mutual Information (MI) between the neural response (R) and the stimulus category (S) quantifies the reduction in uncertainty about one variable given knowledge of the other. Formally, MI can be written in several equivalent forms:

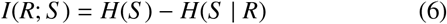

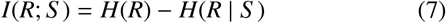

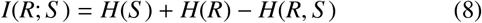

Where *H*(*X*) denotes the marginal entropy of variable X, *H*(*X* | *Y*) the conditional entropy, and *H*(*X, Y*) the joint entropy. Intuitively, *I*(*R*; *S*) measures how much information (in bits) the neural response conveys about the stimulus category. In the present study, S represents the experimental contrast between grating and noise (or natural images) stimuli, and R corresponds to either the broadband (BB) or narrowband gamma (NBG) univariate response (i.e., single-electrode). Thus, MI provides a measure of effect size that quantifies how strongly neural activity discriminates between stimulus classes on a common information-theoretic scale. A value of 1 bit indicates that the response completely resolves the uncertainty between two equally probable stimulus categories, whereas smaller values indicate partial reductions in uncertainty.

This formulation enables a direct comparison of BB and NBG responses in terms of their relative encoding strength. MI was estimated using the Gaussian Copula Mutual Information (GCMI) framework (Ince et al., 2017), which provides bias-corrected, robust (rank-based) estimates suitable for continuous neural data. Specifically, using all available trials per category contrast, the signal at every time point was permuted 200 times for each electrode, randomly assigning the stimulus class labels each time. The maximum value at each time point was taken, and the 95th percentile of this value was used as the threshold for significance. This method corrects for multiple comparisons and provides a Family-Wise Error Rate (FWER) of 0.05. The electrodes listed in the previous section were used separately to compare orientation versus unfiltered noise and orientation versus filtered noise for the BB and NBG signals.

### Co-information analyses

We quantified co-information (co-I) within signals (single electrodes) and between signals (between pairs of electrodes) using the GCMI toolbox (Ince et al., 2017). For the analyses based on the spectral decoupling method (Figures 3-6), the co-I was calculated on the BB and NBG signals separately, using the electrodes reported in the previous sections. The resulting co-I metric (in bits) quantifies the information content, redundant or synergistic, between the two signals. GCMI is a semi-parametric estimator that first transforms the data to have a Gaussian marginal distribution, then uses a Gaussian parametric estimation method. Data are rank transformed to obtain the marginal empirical cumulative density function, which is transformed to Gaussian values. Co-I is formally computed as:

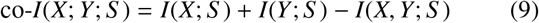

For each time point, *I*(*X*; *S*) corresponds to the mutual information (MI) between the signal at recording site X and stimuli class S. *I*(*Y*; *S*) corresponds to the MI between the signal at recording site Y and stimuli class S. Finally, *I*(*X, Y*; *S*) corresponds to the MI between stimuli class S combining signals from recording sites X and Y.

Positive co-I shows that signals between recording sites contain redundant (or shared) information about the stimulus contrast. Negative co-I corresponds to the synergy between the two variables: the information when considering the two variables jointly is larger than when considering the variables separately. Figure 1D shows a schematic representation of co-I between two signals. It shows the independent information that signal 1 (S1) and signal 2 (S2) (both in white) contain. If there is an overlap in the information represented by the two signals, there is a redundancy (red color) in the information the two responses contain. If the two signals considered together contain more information than could be expected based on the information in the individual signals, there is synergy (blue color). Statistical analyses of co-I charts were performed using a permutation test with 200 permutations and the same maximum statistics method described for the MI analyses, resulting in an FWER of 0.05.

Note that MI and co-I values are reported in units of bits. A value of 1 bit corresponds to a halving of the uncertainty of the trial state when observing the neural response. It is important to keep in mind, though, that these information values are the average per sample. Here, we use a sampling rate of 500 Hz, so a value of 0.1 bits/sample corresponds to an approximate information rate of 50 bits/second (assuming independence) at that specific post-stimulus time point. However, this is just a heuristic comparison; formal estimation of information rates involves more extensive computation to quantify the limit of this information for higher and higher temporal resolution.

### Statistical comparison of co-I, redundancy, and synergy for BB and NBG signals

To statistically compare the BB and NBG signals, we performed two-tailed t-tests separately for co-I, redundancy, and synergy. For within-area contrasts (Figures 3 and 5), t-tests were performed on the normalized count (i.e., percentage) of significant time points (obtained from the previous step described above) per electrode. In the case of the between-area contrasts (Figures 4 and 6), t-tests were performed on the normalized count of significant time points between all pairs of electrodes for the corresponding BB and NBG signals.

**Figure 5:**
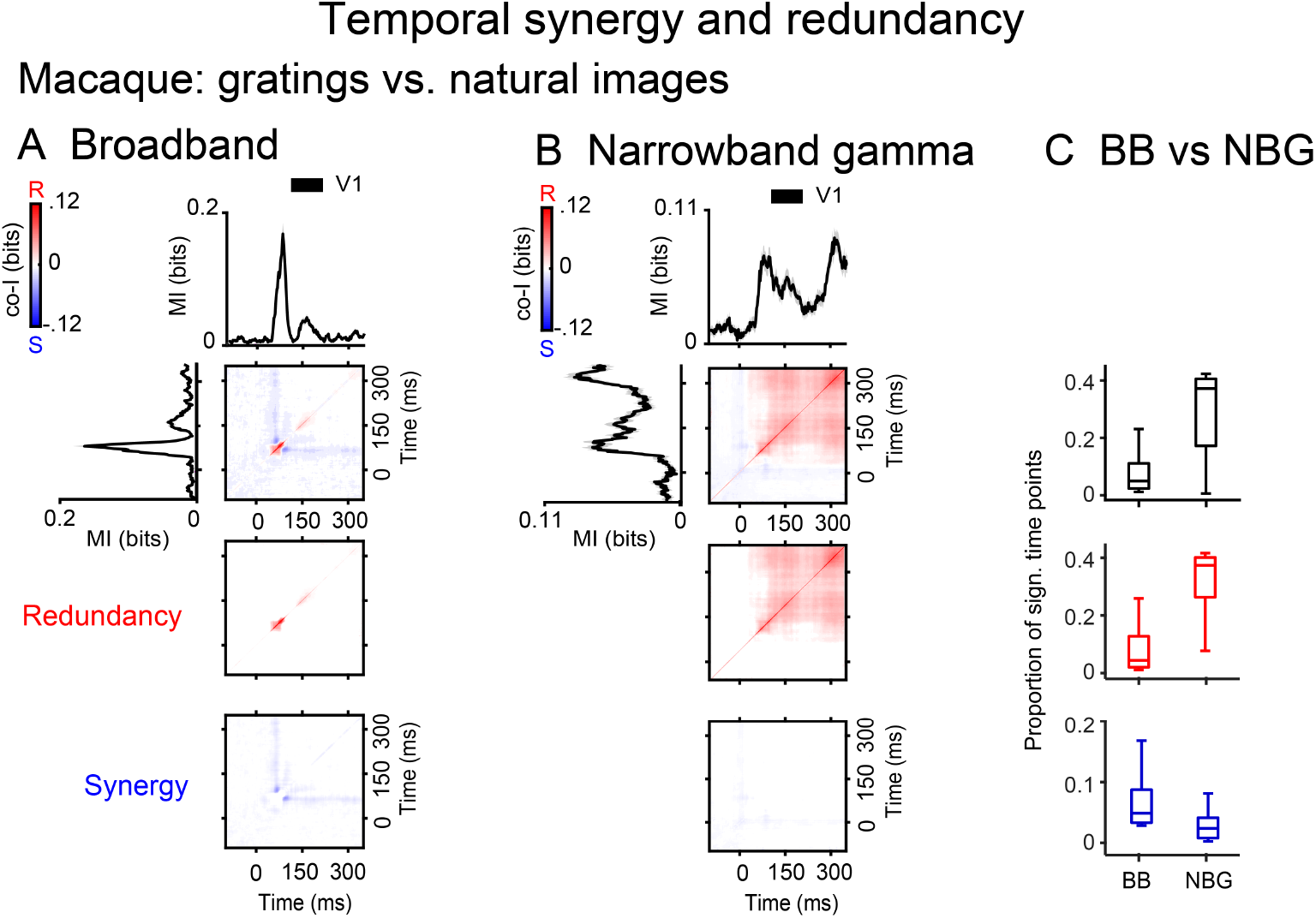
Temporal synergy and redundancy within visual areas in the macaque brain. Co-information revealed synergistic and redundant temporal patterns within the visual cortex in Macaque 1. (**A**) Co-information charts for BB signals in V1. Co-information charts for NBG signals in V1 (**B**). Black traces (V1) represent the MI between gratings and natural images for the corresponding signals. Error bars represent the standard error of the mean (S.E.M) across the corresponding signals. Temporal co-I was computed within visual areas across time points between -100 to 350 ms after image presentation. The average across individual recording sites is shown for the complete co-I chart (red and blue panel); for positive co-I values (redundancy only; red panel); and negative co-I values (synergy only; blue panel). (**C**) Box plots display the sample median alongside the interquartile range. Samples represent the percentage of significant time points observed in each electrode for co-I (black boxes), redundant (positive values; red boxes), and synergistic information (negative values; blue boxes) in the corresponding BB and NBG signals.

**Figure 6:**
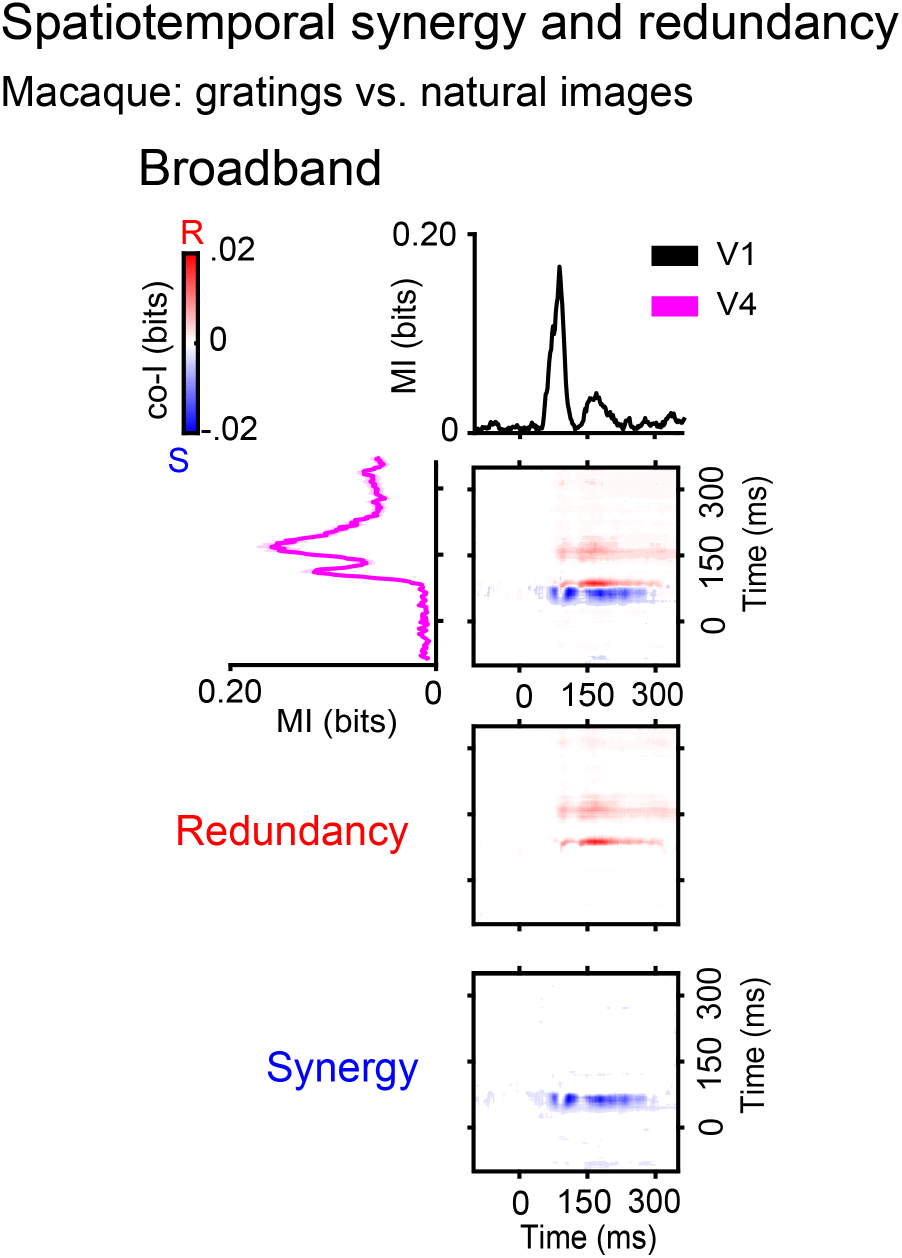
Spatiotemporal synergy and redundancy between visual areas in the macaque brain. Co-information revealed synergistic and redundant spatiotemporal patterns between BB signals across recording sites. Black traces (electrodes in V1) and magenta traces (electrodes in V4) represent the MI between gratings and noise trials. Error bars represent the standard error of the mean (S.E.M) between the corresponding electrodes. Co-I was computed between each pair of electrodes (V1-V4) and across time points between -100 to 350 ms after image presentation. The average across inter-areal electrode pairs is shown for the complete co-I chart (red and blue panel); for the positive co-I values (redundancy only; red panel); and the negative co-I values (synergy only; blue panel).

Let 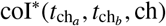 represent the significant co-information, where 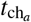 and 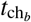 denote time indices, and ch denotes the channel index. The count of significant co-information over the first two dimensions (time × time) for each channel ch is given by:

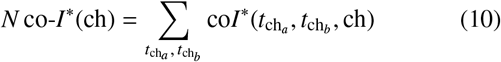

where 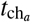 and 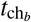 range over all time indices.

### Eye movement recordings and microsaccade detection

Eye movements and pupil size were recorded binocularly at 1000 Hz using an Eyelink 1000 system (SR Research) with infrared illumination. Eye signals were calibrated before each recording session using a standardized fixation task. Microsaccades were detected using an unsupervised clustering method described by Otero-Millan et al. (2014). Candidate microsaccades were identified as periods during which eye velocity exceeded 5 degrees/sec. Saccade onset and offset were defined as the first and last points, respectively, where the velocity or acceleration exceeded the detection thresholds surrounding the peak velocity. We excluded microsaccades occurring during blinks or inter-trial intervals, based on eyelink pupil data and blink events. Overlapping saccade candidates were merged, and a minimum inter-saccadic interval of 30 ms was enforced. For each detected microsaccade, we calculated metrics including amplitude (Euclidean distance between the start and end positions), direction (angle of the amplitude vector), peak velocity, and microsaccade duration. Microsaccade frequency (Hz) was calculated as event count per trial duration.

---

**Figure S1:**
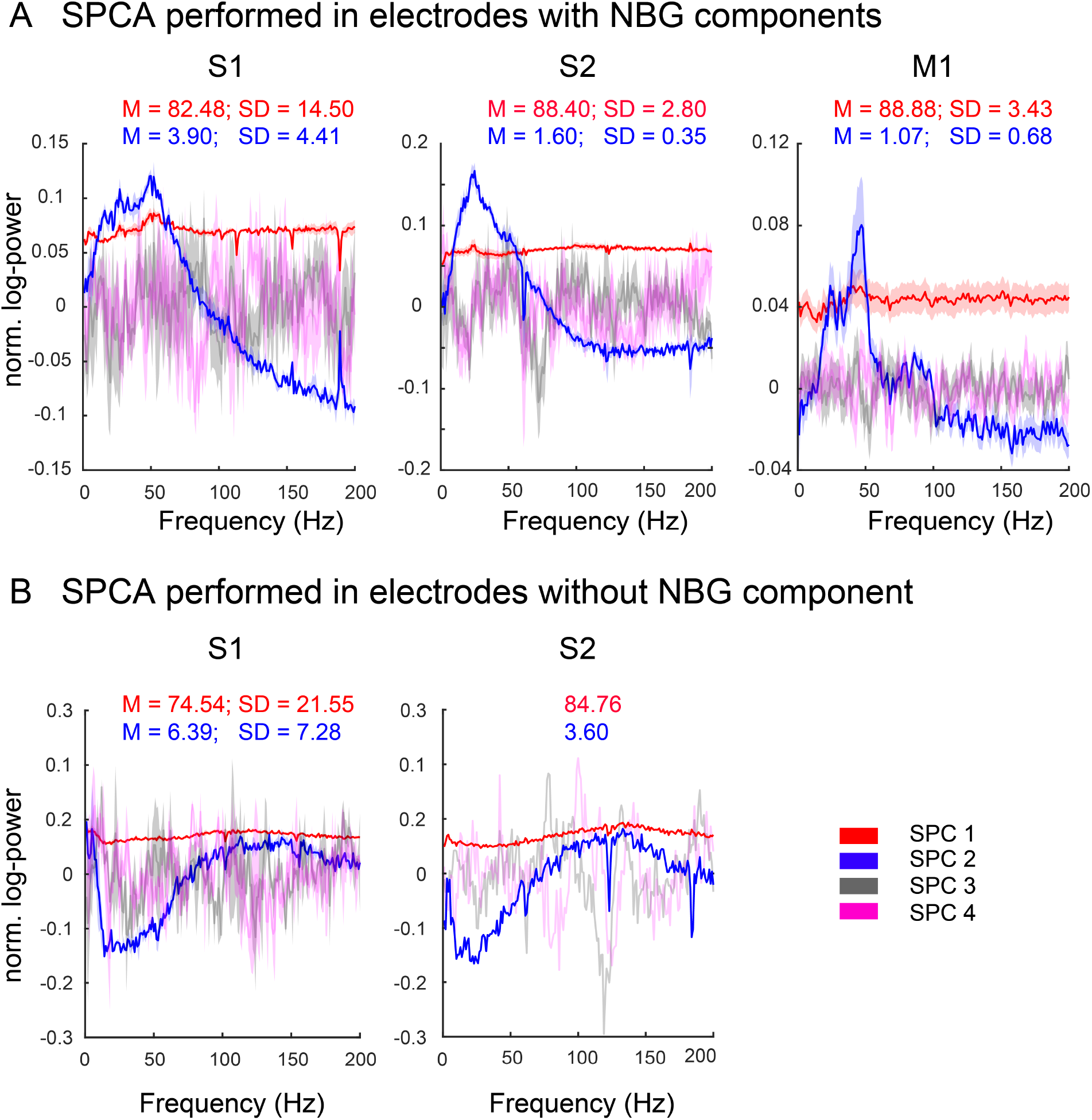
SPCA component for S1, S2, and M1. **(A)** Averaged traces showing a characteristic NBG component for S1 (4 out of 7 electrodes), S2 (5 out of 6 electrodes), and M1 (63 out of 63 electrodes). While SPC1 (red) does not show a characteristic peak across frequencies (BB component), SPCA2 (blue) shows increased magnitude spanning the gamma band (NBG). The mean and standard deviation of the variance explained by the SPC1 and SPC2 components across channels are depicted in red and blue, respectively. Unlike SPC1 and SPC2, SPC3 (gray) and SPC4 (magenta) exhibit high channel variability without a characteristic peak. **(B)** Averaged traces for electrodes without a characteristic NBG component for S1 (2 out of 7 electrodes), S2 (1 out of 6 electrodes). The mean and standard deviation of the variance explained by the SPC1 and SPC2 components across channels are depicted in red and blue, respectively. In contrast, SPC1 shows a broadband component, and SPC2 mainly peaks in the lower-frequency range (<10 Hz).

**Figure S2:**
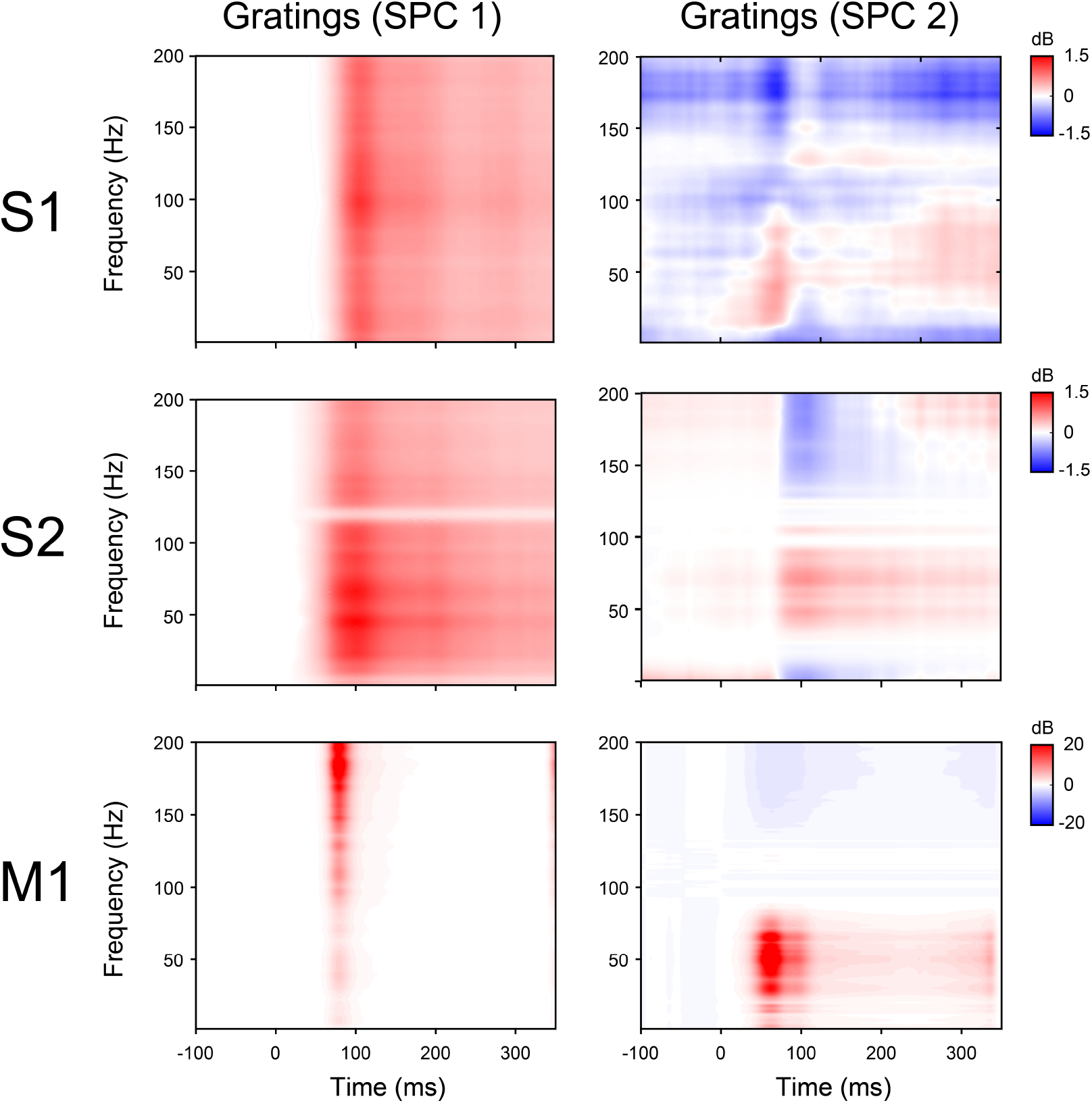
Time–frequency spectral reconstruction using SPC1 and SPC2 in S1, S2, and M1. For each electrode and trial with grating stimuli, Welch power spectra were computed with a single full-length segment (nfft = 500), retaining positive-frequency bins up to 200 Hz. To capture spectral structure, spectra were log-normalized at each frequency, and spectral PCA (SPCA) was performed on the frequency–frequency covariance matrix. The first two components (SPC1 and SPC2) were treated as fixed spectral motifs. We then computed an SPC-specific time course: the signal was filtered with a bank of complex Morlet wavelets (5 cycles per frequency), the squared magnitude was taken to obtain band-power over time, convolution edge samples were discarded, each band’s power trace was normalized to unit mean, a natural-log transform was applied, and the resulting log-power matrix was projected onto the SPC vector to yield a single time course for that component. To reconstruct absolute time–frequency (TF) power, we (1) paired this time course with the component’s frequency profile to form a separable log-power map, (2) exponentiated that map to convert it into a multiplicative power modulation, and (3) multiplied, at each frequency, by the electrode mean Welch spectrum to return to absolute power units. We then baseline-normalized per frequency to -100 to 0 ms and reported the values in dB (10 log10 of the baseline ratio). Panels show TF maps averaged across channels. Power is expressed in decibels (dB) relative to the baseline period. While SPC1 spans most of the frequency axis, SPC2 shows non-zero power predominantly in the gamma range (30-80 Hz).

**Figure S3:**
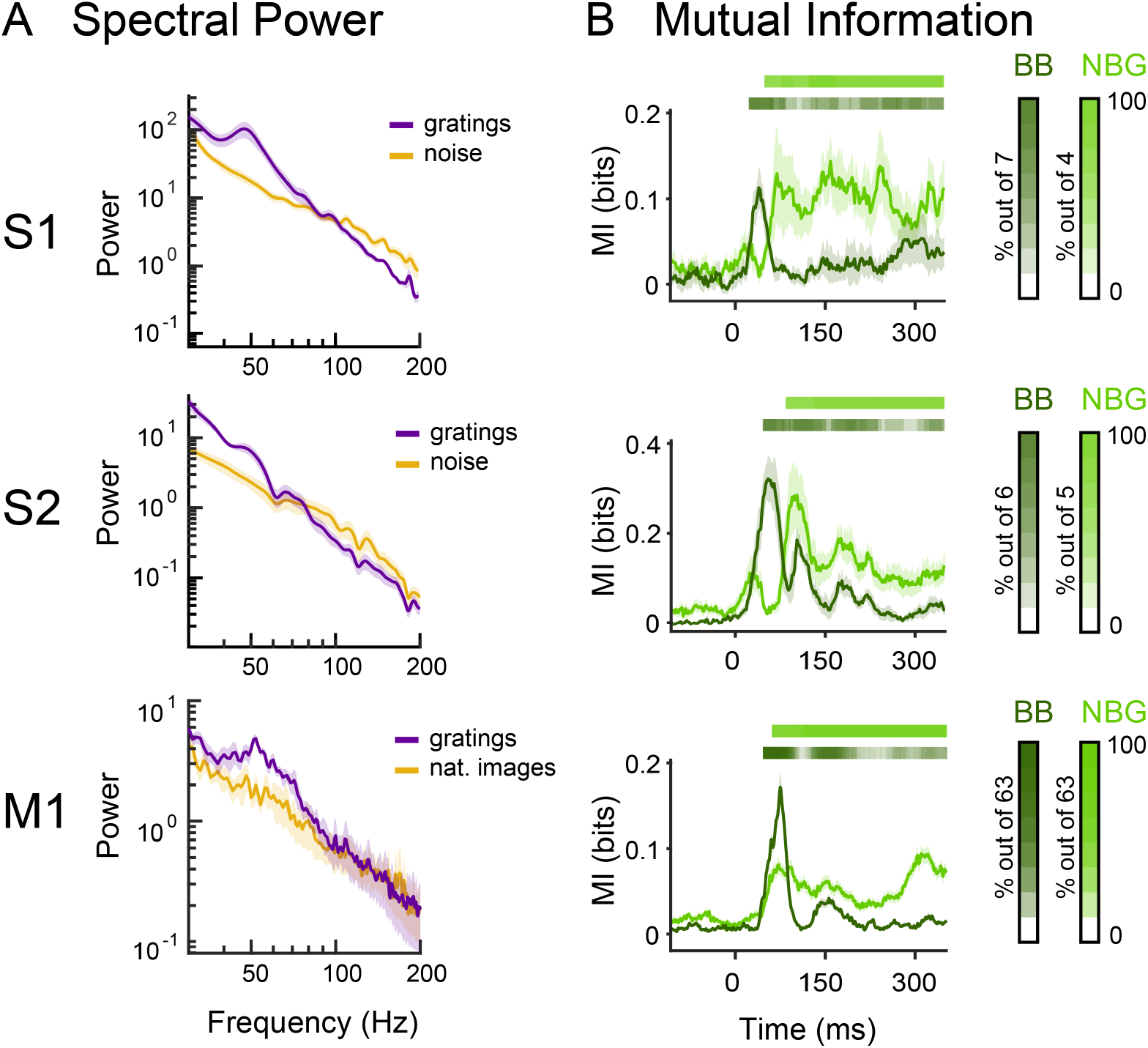
Spectral power and mutual information for S1, S2, and M1. **(A)** Power spectral density for gratings (purple) and noise (yellow) stimuli for S1 and S2, and for gratings and natural images (yellow) for M1. Error bars represents the standard error of the mean (S.E.M) across electrodes (S1: V1,V2; S2: V1, V2, V3, and M1: V1). **(B)** Mutual information analyses between gratings and noise images for the BB component for S1 and S2 (dark green), and between gratings and natural images for M1 (dark green). Mutual information between gratings and noise images for the NBG component for S1 and S2 (light green), and between gratings and natural images for M1 (light green). The shaded horizontal bars represent significant MI values at the single-electrode level, with the colors representing the percentage of electrodes showing a significant effect per time point for BB (dark green) and NBG (light green). Error bars represent the S.E.M of the MI values across electrodes.

**Figure S4:**
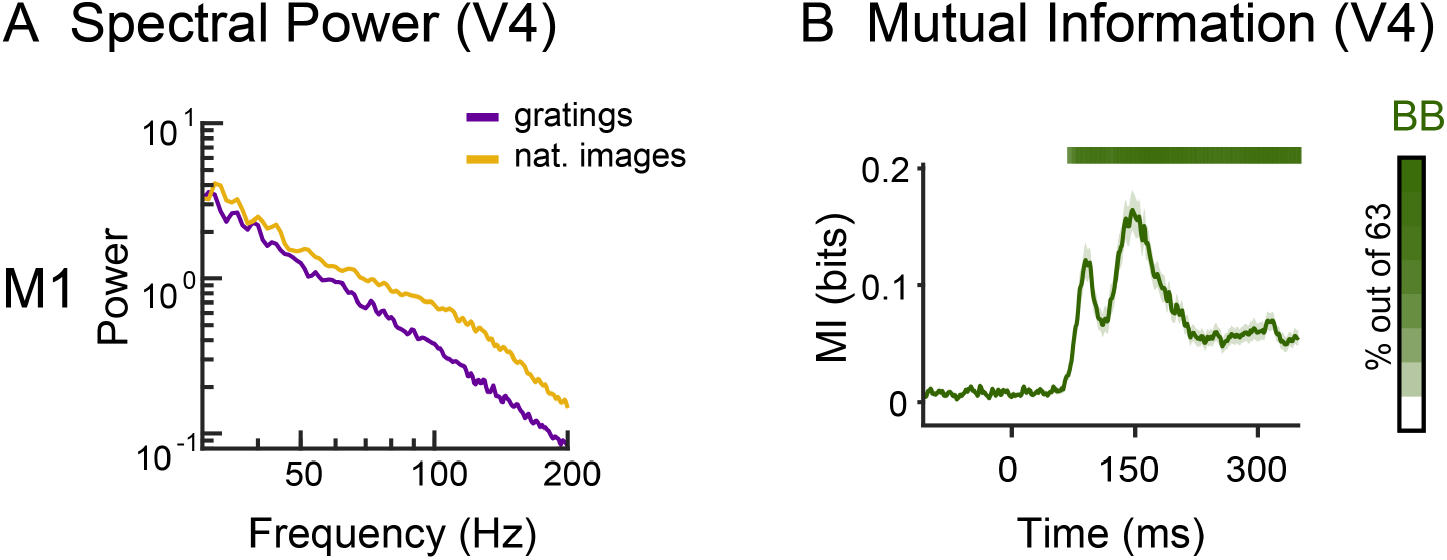
Spectral power and mutual information for M1 in area V4. **(A)** Power spectral density for gratings (purple) and noise and natural images (yellow). Error bars represent the standard error of the mean (S.E.M) across electrodes in V4. **(B)** Mutual information analyses between gratings and natural images for M1 (dark green). The shaded horizontal bars represent significant MI values at the single-electrode level, with the colors representing the percentage of electrodes showing a significant effect per time point for BB (dark green). Error bars represent S.E.M of the MI values across electrodes.

**Figure S5:**
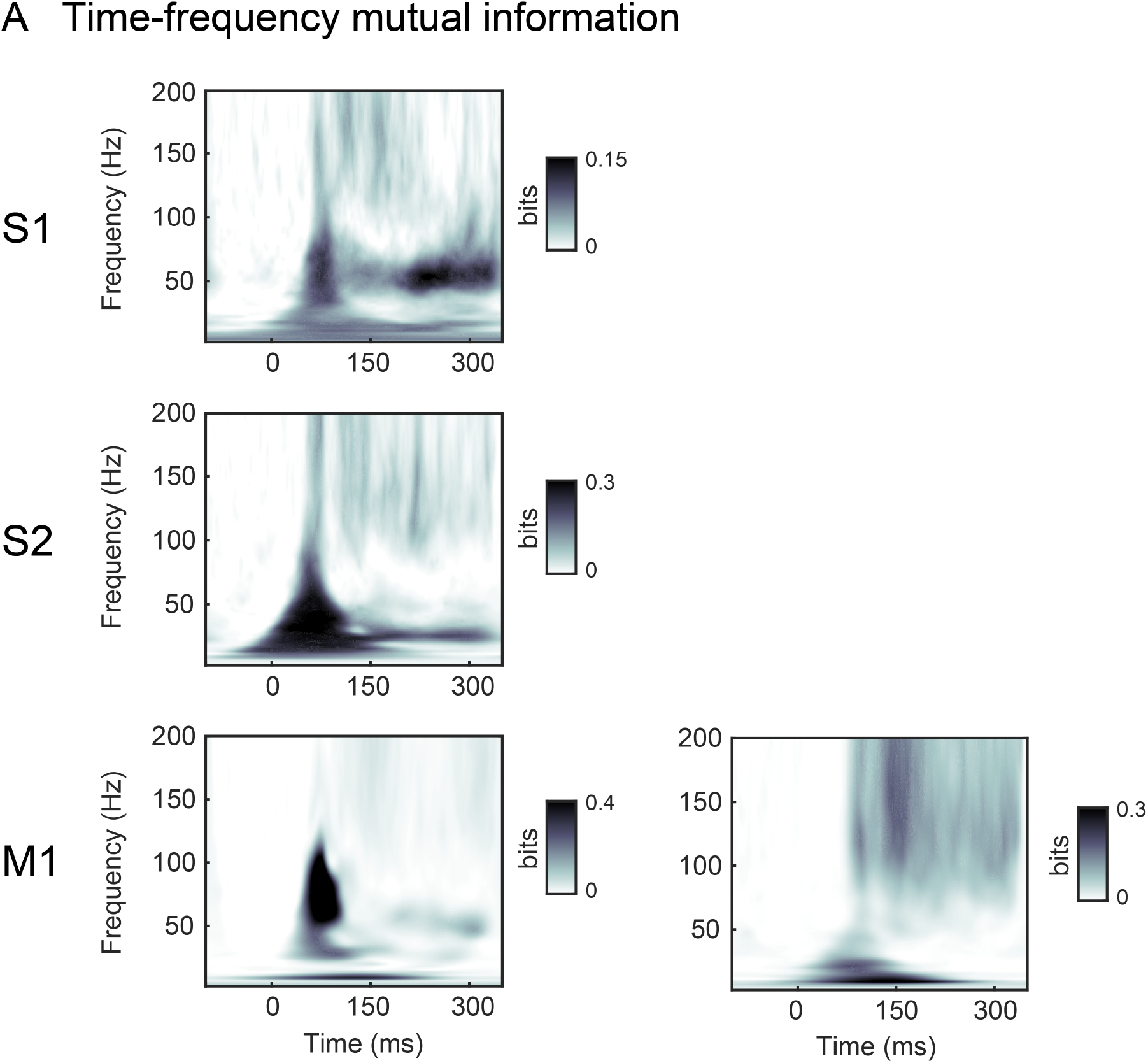
Time-frequency mutual information for S1, S2, and M1. **(A)** Mutual information (MI) between gratings and noise for S1 and S2, and between gratings and natural images for M1. MI was computed from power values across trials for each time-frequency bin from 1 to 200 Hz, within the -100 to 350 ms window around stimulus onset. MI represents the effect size (in bits) of the difference in spectral power for each corresponding stimulus comparison. Plots represent the average MI across electrodes (S1: 7 electrodes [V1 and V2 area]; S2: 6 electrodes [V1, V2 and V3 area], M1: 63 electrodes in V1 (left panel), and 63 electrodes in V4, right panel).

**Figure S6:**
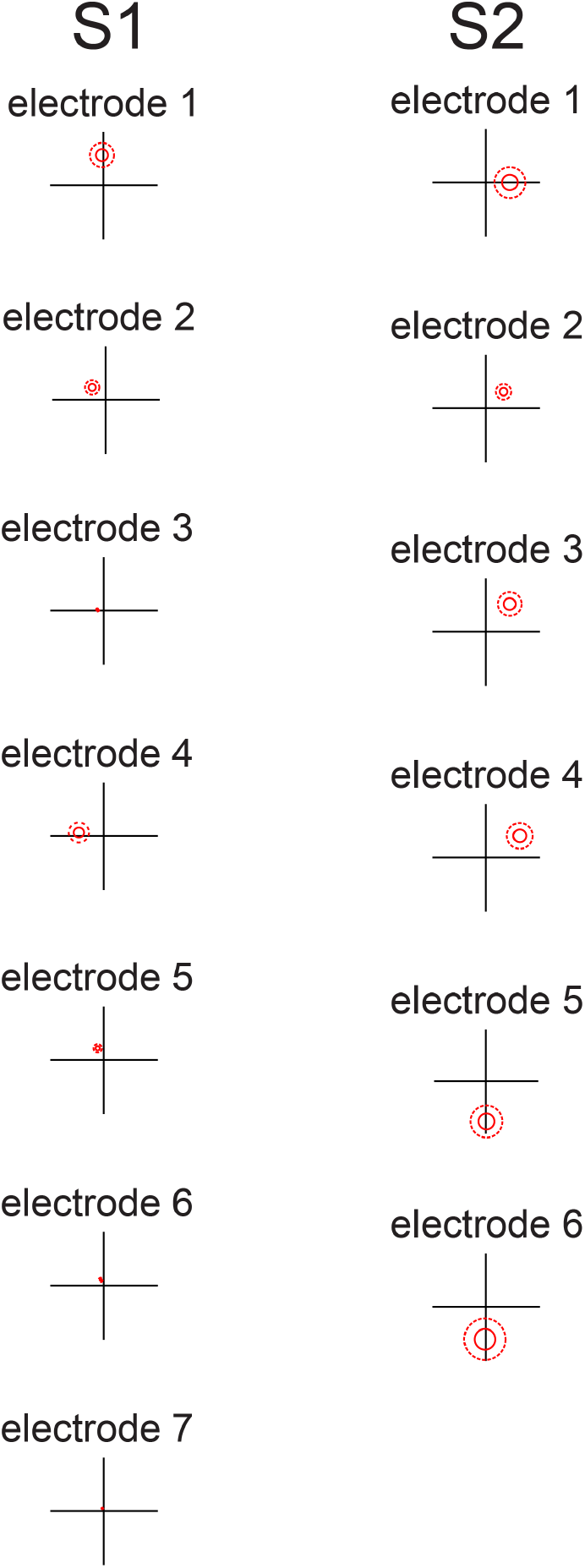
location of population receptive fields (pRF) for S1 and S2. The population receptive field for each electrode was defined by a Gaussian, indicated by the 1- and 2-sd contours (solid and dotted red lines).

**Figure S7:**
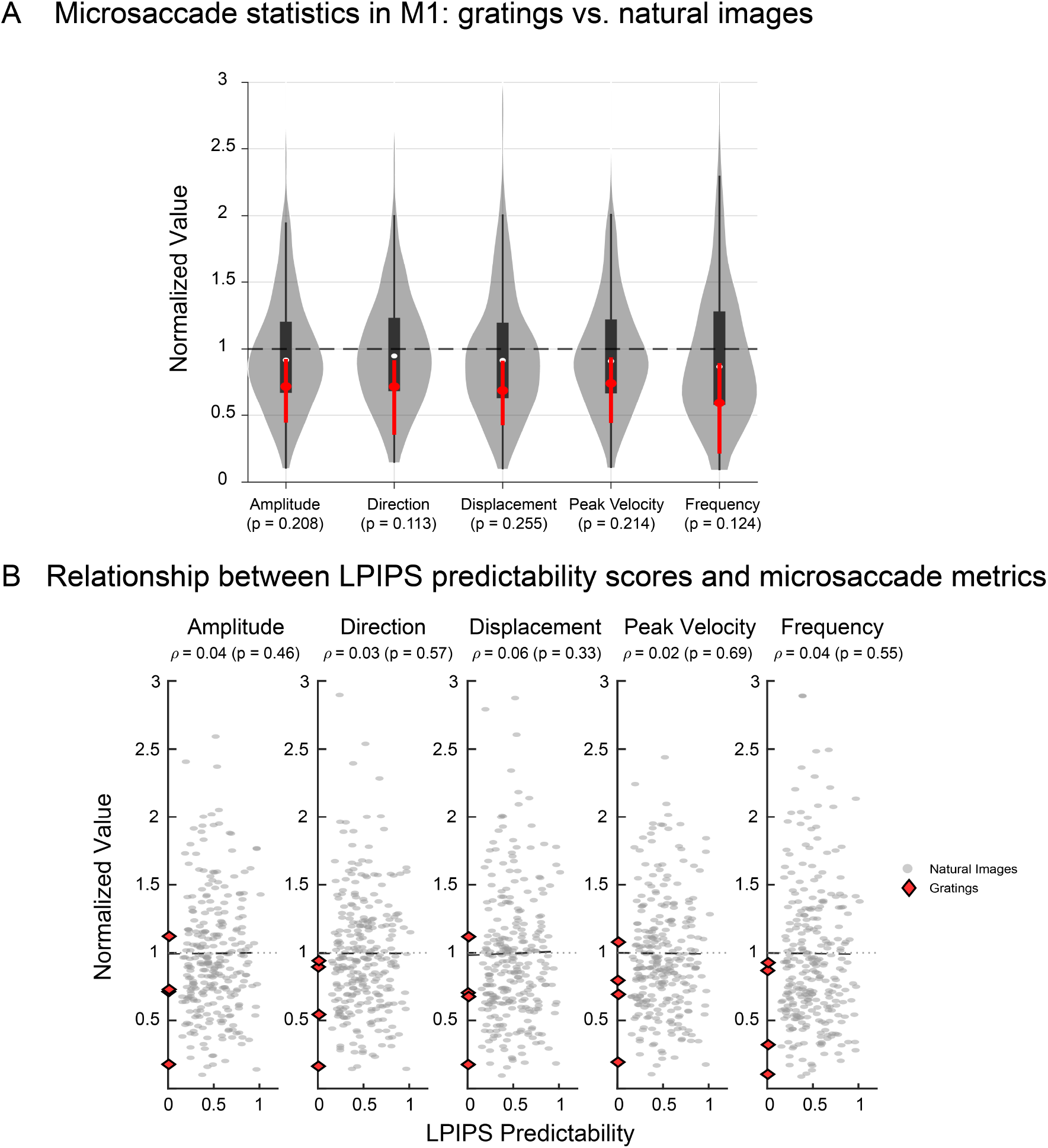
Eye Movement Analysis. To address potential oculomotor confounds, we analyzed eye-tracking data collected during the presentation of both natural images and gratings, which were used in all other analyses. **(A)** Microsaccades were detected using the algorithm as described in Otero et al. (2014). We extracted five kinematic metrics for comparison: amplitude, direction, displacement, peak velocity, and frequency. For each metric, we compared the distributions of the natural images and gratings. For visualization, all metric values were normalized to the mean of the Natural Image condition (Natural images Mean = 1). We assessed statistical significance using the nonparametric Wilcoxon rank-sum test. In the resulting plots, the natural image distribution is shown as a violin plot (gray), while the gratings are summarized by their median (red circle) and interquartile range (red vertical line). **(B)** The relationship between LPIPS predictability scores and microsaccade metrics. Grating stimuli are shown in red, and natural images are shown in gray. Grating stimuli follow the same distribution as natural images; they do not form an outlier cluster that drives a spurious trend. Across the pooled dataset, we found no significant correlation between image statistics and microsaccade dynamics (Spearman correlation, p > 0.05 for all metrics).

